# Chromatin-contact atlas reveals disorder-mediated protein interactions and moonlighting chromatin-associated RBPs

**DOI:** 10.1101/2020.07.13.200212

**Authors:** Mahmoud-Reza Rafiee, Julian A Zagalak, Sviatoslav Sidorov, Sebastian Steinhauser, Karen Davey, Jernej Ule, Nicholas M Luscombe

## Abstract

RNA-binding proteins (RBPs) play diverse roles in regulating co-transcriptional RNA-processing and chromatin functions, but our knowledge of the repertoire of chromatin-associated RBPs (caRBPs) and their interactions with chromatin remains limited. Here, we developed SPACE (Silica Particle Assisted Chromatin Enrichment) to isolate global and regional chromatin components with high specificity and sensitivity, and SPACEmap to identify the chromatin-contact regions in proteins. Applied to mouse embryonic stem cells, SPACE identified 1,459 chromatin-associated proteins, ∼48% of which are annotated as RBPs, indicating their dual roles in chromatin and RNA-binding. Additionally, SPACEmap stringently verified chromatin-binding of 404 RBPs and identified their chromatin-contact regions. Notably, SPACEmap showed that about half of the caRBPs bind to chromatin by intrinsically disordered regions (IDRs). Studying SPACE and total proteome dynamics from mES cells grown in 2iL and serum medium indicates significant correlation (R = 0.62). One of the most dynamic caRBPs is Dazl, which we find co-localized with PRC2 at transcription start sites of genes that are distinct from Dazl mRNA binding. Dazl and other PRC2-colocalised caRBPs are rich in intrinsically disordered regions (IDRs), which could contribute to the formation and regulation of phase-separated PRC condensates. Together, our approach provides an unprecedented insight into IDR-mediated interactions and caRBPs with moonlighting functions in native chromatin.

## Introduction

RBPs participate in regulating transcription as well as other aspects of co-transcriptional RNA regulation (1,2). Indeed, it is known that transcriptional and post-transcriptional processes are integrated to coordinate alternative splicing and polyadenylation (3,4), RNA stability (5,6) and subsequent translation in the cytoplasm (7). Furthermore, RBPs promote biomolecular condensate formation, and were reported to contribute to the functionality of enhancers, transcription factors and RNA Pol II (8-10). **Considering all these potential RBP-chromatin interactions, the question is which RBPs join the repertoire of chromatin-associated proteins**. This is particularly important as changes in the dynamics of RBPs are generally implicated in cancer and neurodegenerative diseases (11,12).

Global UV-crosslinkable RNA interactome capture based on oligo-dT capture, click chemistry or organic phase separation have identified over ∼2,300 candidate RBPs (13). However, these methods are not able to distinguish those RBPs that associate with chromatin (chromatin-associated RBPs, caRBPs). ChIP-seq has been used to assess the association of dozens of RBPs with chromatin (2,14), but its application is limited by the availability and specificity of antibodies. Thus, **methods are needed that provide a global view of caRBPs** with high specificity and throughput.

Traditionally, chromatin is isolated by cellular fractionation and precipitation (15). However, the results are ambiguous due to the abundant cytoplasmic contaminations that remain in the nuclear fraction and precipitate together with chromatin. In order to enhance specificity, DNA-labelling by ethynyl deoxy-uridine (EdU) was implemented to isolate chromatin fragments by click-chemistry and streptavidin beads (16,17). However, incorporation of modified nucleotides into DNA can’t preserve the natural conditions of chromatin. **Additionally, current chromatome methods are unable to determine the chromatin-protein contact sites, which is essential to reliably understand how proteins are integrated to the chromatin network**.

Here, we present SPACE (Silica Particle Assisted Chromatin Enrichment), a straightforward and highly sensitive method that relies on silica magnetic beads for chromatin purification. To demonstrate the power of the method, we evaluated SPACE by studying the global chromatin composition of mES cells. We successfully identified previously reported DNA- and chromatin-binding proteins, as well as many caRBPs. **Surprisingly, RBPs comprise ∼48% of the proteins obtained from the chromatome**. To understand how RBPs bind to chromatin, we developed SPACEmap. We found that intrinsically disordered regions (IDRs) are frequently employed by chromatin proteins, including caRBPs, for chromatin-binding. Taken together, we demonstrate that the various applications of SPACE provide flexible, highly sensitive and accurate approaches for studying dynamics of chromatin-associated proteins, which has proven particularly valuable to expand the knowledge of RBP-chromatin interactions.

## Material & Methods

### Mass spectrometry and proteomics data analysis

The details of sample preparation using SPACE, SPACE-SICAP and ChIP-SPACE procedures are provided in the Supplementary Material. Briefly, the cells were crosslinked using formaldehyde 1% final concentration in the medium of the cells (v/v) within 10 min. Then the cells were washed with PBS, and frozen. After the SPACE process (described in Supplementary Material), the proteins were digested on the beads using trypsin and LysC. Following sample preparation, peptides were separated on a 50□cm, 75□µm I.D. Pepmap column over a 120□min gradient for SPACE and SPACE-SICAP, or a 70min gradient for ChIP-SPACE. Peptides were then injected into the mass spectrometer (Orbitrap Fusion Lumos) running with a universal Thermo Scientific HCD-IT method. Xcalibur software was used to control the data acquisition. The instrument was run in data-dependent acquisition mode with the most abundant peptides selected for MS/MS by HCD fragmentation. RAW data were processed with MaxQuant (1.6.2.6) using default settings. MSMS spectra were searched against the UniProt (Swissprot) database (*Mus musculus*) and database of contaminants. Trypsin/P and LysC were chosen as enzyme specificity, allowing a maximum of two missed cleavages. Cysteine carbamidomethylation was chosen as the fixed modification, and methionine oxidation and protein N-terminal acetylation were used as variable modifications. Global false discovery rate for both protein and peptides was set to 1%. The match-from-and-to and re-quantify options were enabled, and Intensity-based quantification options (iBAQ) were calculated.

### Quantitative proteomics, statistical and computational analysis

The protein groups were processed in RStudio using R version 4.0.0. The proteins only identified by site, Reverse and potential contaminants were filtered out. For all datasets in this study Gene Ontology (GO) and other information were downloaded from UniProt and DAVID Gene Ontology database. For the SPACE experiments (related to Figure 2-3), the crosslinked samples were compared with non-crosslinked samples by SILAC ratios calculated using MaxQuant. In total we did 2 forward (heavy SILAC is crosslinked and light SILAC is not crosslinked) and 6 reverse experiments (light SILAC is crosslinked and heavy SILAC is not crosslinked). We considered proteins identified using at least 1 forward and 1 reverse experiments (> 2 assays in total) for statistical analysis. Bayesian moderated t-test p-values and Benjamini-Hochberg (BH) adjusted p-values (adj. p-value) were calculated by limma package (18). The limma package calculated fold-changes (FC) as follows: log2FC = mean(log2(crosslinked/non-crosslinked)). We, therefore, considered log2FC > 1 and adj. p-value <0.01 as highly significant, and log2FC > 1 and adj. p-value <0.1 as significantly enriched proteins using SPACE. The SPACE experiments were carried out using varying cell numbers. We used 2.5 million cells for forward SILAC labelling experiments. We also used 500,000, 100,000 and 20,000 cells for reverse SILAC labelling experiments (related to Figure 2A and Supplementary Figure 2F). We also performed statistical analysis using proteins identified by 2 out of 2 replicates for each cell number. The statistical thresholds were applied as described above to identify the enriched proteins.

We obtained Gene Ontology (GO) data from UniProtand DAVID databases and merged them with our protein datasets. We categorized the enriched proteins to potential true positive (PTP) if they are involved in a function or a biological process that is related to chromatin. Otherwise, we considered them as potential false positive (PFP) groups (related to Figure 2C and Supplementary Figure 2A). To categorize the proteins we searched for specific keywords in the following columns: Gene.ontology.molecular.function, Gene.ontology.biological.process, Keywords, GOTERM_MF_DIRECT and GOTERM_BP_DIRECT. To determine PTPs, we looked for the following keywords: 1-chromatin remodelling 2-chromatin modification, chromatin modifier, nucleosome assembly, chromatin silencing, histone methylation, chromatin assembly 3-DNA replication, DNA repair, double-strand break repair, DNA damage response, DNA helicase, DNA/RNA helicase, response to DNA damage, telomere maintenance 4-chromosome segregation, chromosome separation, condensed chromosome kinetochore, sister chromatid biorientation 5-transcription factor activity, transcription co-activator, transcription coactivator, transcription corepressor, transcriptional activator, transcriptional repressor or transcription factor binding 6-regulation of DNA-templated transcription, transcription DNA-templated, regulation of transcription, transcription by RNA polymerase II, transcription regulation, RNA polymerase II transcription, transcription from RNA polymerase II, transcription-dependent tethering of RNA polymerase II, rRNA transcription 7-posttranscription, post-transcription 8-cell cycle 9-splicing, spliceosome 10-RNA processing, rRNA processing, polyadenylation, RNA 3’-end, RNA cleavage, RNA modification, RNA capping, RNA modification 11-RNA transport, and we grouped the proteins respectively. Among the rest of the proteins, we looked for the following keywords: 1-translation, 2-metabolic process, oxidase, 3-cell adhesion, 4-protein folding, and 5-protein transport, and we grouped the proteins respectively. If these proteins are not present in nucleus, we considered them as PFPs. If they are present in nucleus, we considered them as neither PFP nor PTP. Because they might have uncharacterised chromatin-related functions in nucleus. Proteins that do not contain any of these keywords in their GO columns were considered as “miscellaneous”. Miscellaneous proteins that are known DNA/chromatin-binders were considered as PTPs. Miscellaneous proteins that are not present in the nucleus were considered as PFPs. The miscellaneous proteins that are present in nucleus were considered as neither PTP nor PFP, because they might have uncharacterised chromatin-related functions in nucleus.

The enriched proteins were also categorized to known DNA/chromatin-binders and proteins that are “present in the nucleus” (but not DNA/chromatin-binders). The rest of the proteins were considered as “unexpected”. Specifically, we looked for “DNA-binding, DNA binding, DNA_BIND” keyword in the following columns to determine known DNA-binders: Gene.ontology.molecular.function, DNA.binding, GOTERM_MF. We also looked for “chromatin” and “chromosome” keywords in the following columns to determine known chromatin/chromosome-binding proteins: GO.molecular.function, GO.cellular.component, Subcellular.location.CC, GOTERM_MF. We looked for the “nucleus, nucleolus, nucleoplasm, nuclear matrix, nuclear envelope” keywords in the following columns to determine proteins present in the nucleus: GO.cellular.component., Subcellular.location..CC., GOTERM_CC. To determine RNA-binding proteins, we looked for “RNA-binding/UTR-binding” in the following columns: GO.molecular.function and GOTERM_MF.

SPACE-SICAP (related to Figure 2-3) was carried out using 5 replicates. Proteins identified using at least 2 replicates were considered for statistical analysis. The crosslinked samples were compared with the non-crosslinked samples by SILAC iBAQ values. The crosslinked samples and non-crosslinked samples were normalized separately using quantile-normalization from preprocessCore package (https://github.com/bmbolstad/preprocessCore). If maximum 2 out of 5 replicates had no values (missing values), they were imputed using the mean of the other replicates. If all 5 replicates in the non-crosslinked samples were missing, minimum iBAQ values were used for the imputation. Bayesian moderated t-test p-values and BH adj. p-values were calculated by limma package. We considered log2FC > 1 and adj. p-value <0.01 as highly significant, and log2FC > 1 and adj. p-value <0.1 as significantly enriched proteins using SPACE-SICAP.

SPACE and SPACE-SICAP results were compared with total proteome (19), DmChP (17) and chromatin pelleting (20). Published data were downloaded and re-analysed using MaxQuant. All the datasets were produced using mES cells grown in 2iL medium. DmChP dataset contains 8 EdU-plus experiments, and 7 EdU-minus experiments. For the sake of consistency among the datasets, we re-analysed DmChP data using MaxQuant label-free quantification by iBAQ values. We filtered proteins identified using at least 2 EdU-plus experiments for statistical analysis using limma package. The crosslinked samples and non-crosslinked samples were normalized separately using quantile-normalization. If all 7 EdU-minus replicates were missing, they were imputed with minimum intensities. We considered log2FC > 1 and adj. p-value <0.01 as highly significant, and log2FC > 1 and adj. p-value <0.1 as significantly enriched proteins using DmChP. Chromatin pelleting dataset contains 3 replicates, and intrinsically doesn’t have a negative control. Thus, proteins identified with at least 2 replicates were considered for the comparisons with the other datasets. The proteins were categorized as described previously to known “DNA/chromatin-binders”, “present in the nucleus” and “unexpected” proteins. Fisher’s exact test was used to show statistically significant differences between the datasets with *** for p-value < 0.001, ** for p-value ≤ 0.01 and * for p-value ≤ 0.05.

For the SPACEmap experiment (related to Figure 4), the crosslinked fraction was compared with the released fraction by peptide intensities using 3 replicates for each fraction. The samples were normalized using quantile-normalization from preprocessCore package. If all 3 replicates of the released fraction or the crosslinked fractions were missing, they were imputed with minimum intensities. If 1 out of 3 replicate was missing, it was imputed with the mean of the other two replicates. Moderated t-test p-values and BH adj. p-values were calculated by limma package. Log2(crosslinked/released) > 0.4 and adj. p-value < 0.1 were considered as differentially enriched peptides.

For the comparative SPACE experiment and total proteome analysis (related to Figure 5), the 2iL (heavy SILAC) samples were compared with serum samples (light SILAC) by ratios calculated using MaxQuant. Moderated t-test and BH adj. p-values were calculated by limma package. Log2(2iL/serum) >1 and adj. p-value <0.1 were considered as significantly enriched proteins. Interaction network determined only by experiments was downloaded from String database and visualized by Cytoscape 3.8.

For the Dazl ChIP-SPACE experiment (related to Figure 6), the RNase-treated and non-treated samples were compared by label-free iBAQ values using 3 replicates for each condition. Moderated t-test p-values and BH adj. p-values were calculated by limma package. Log2(RNase-untreated/treated) > 1 and adj. p-value <0.1 were considered as differentially enriched proteins.

### Dazl ChIP-seq experiment and data analysis

Details of the ChIP procedure and data analysis were described in Supplementary Material. Briefly, mES cells were grown in 2iL medium. The cells were detached and fixed by 1.5% formaldehyde in PBS for 15min. Chromatin was solubilized by sonication and sheared to < 500 bp fragments, with the peaks about 200-300 bp. Dazl immunoprecipitation was carried out using CST antibody #8042 overnight at 4 °C. Following washing steps, chromatin was eluted, and DNA was purified by SPRI beads. Library was prepared for the Illumina platform. Sequencing was carried out using 100nt reads on paired-end mode by HiSeq4000. Reads were trimmed, aligned to the mouse genome (mm10) using Bowtie2, and duplicated reads were removed with samtools. The ChIP quality was evaluated by cross-correlation using the “SPP” tool as suggested by ENCODE ChIP-seq guidelines. Peak calling was performed using MACS2. Reproducibility of the ChIP replicates and final peak selection was assessed using the IDR pipeline at a 1% IDR cutoff for the final list of the peaks. Dazl peaks annotation into genomic features was done using ChIPseeker R package with 3kb around TSS set for promoter region window. The ChIP-seq profiles of Suz12, Aebp2 and H3K27me3 were obtained from published data (21), and were compared with Dazl ChIP-seq by deepTools 2.

### Dazl iCLIP and data analysis

The iCLIP assay was carried out as previously described (22). Briefly, mESCs were grown in 2iL medium. Cells were UV cross linked, lysed and IP performed using 1:70 DAZL antibody (CST #8042) in IP. RNaseI was used at 0.4U/mg cell lysate per IP. Finalised libraries were sequenced as single end 100bp reads on Illumina HiSeq 4000. Processing of DAZL iCLIP raw data was carried out using iMaps (https://imaps.genialis.com/). The demultiplexed and quality-controlled data was mapped to mm10 genome assembly using STAR (2.6.0) with default settings. Both PCR duplicates and reads that did not map uniquely to the genome were discarded.

### Cell culture

The 46C mES cells were cultured using either 2i + LIF (2iL) medium or standard mESC serum medium + LIF. The 2iL medium consists of DMEM:F12 for SILAC, Glutamax, N2 supplement, non-essential amino acids, B27 supplement, β-mercaptoethanol (all from Gibco), CHIR99021 3uM (Sigma-Aldrich), PD0325901 1uM (Sigma-Aldrich) and LIF 100 ng/ul (proteintech). The 2iL medium represents the ground-state of the mouse ES cells while serum state represents the meta-stable state. To label the cells with heavy amino acids, ^13^C_6_ ^15^N_4_ L-Arginine and ^13^C_6_ ^15^N_2_ L-Lysine were added to the 2iL medium. To label the cells with light amino acids, ^12^C_6_ ^14^N_4_ L-Arginine and ^12^C_6_ ^14^N_2_ L-Lysine were added to the medium.

### Domain analysis

For details of domain analysis please see Supplementary Material and Supplementary Figure 4F-H. Briefly, we searched domains and intrinsically disordered regions (IDRs) in the proteins from the crosslinked and released SPACEmap fractions using InterProScan v5.47-82.0. We excluded matches that did not represent domains or IDRs and merged highly overlapping retained matches to obtain consensus matches for further analysis. Next, we searched domains and IDRs that matched peptides from the crosslinked and released SPACEmap fractions. We postulated that a domain or an IDR matched a peptide if it overlapped with the peptide or resided no farther than 10 amino acids from the ends of the peptide. Because some domains are rich in arginine and lysine residues. As a result, tryptic peptides are too short for mass spectrometry. Finally, we clustered domains that were matched by peptides from the crosslinked fraction to obtain more general domain types.

## Results

### Designing SPACE and related methods to enrich for chromatin-associated proteins

Silica matrices (columns or beads) are widely used to purify DNA in diverse contexts, but they have not been applied to chromatin purification yet. We reasoned that some regions of DNA are likely to remain accessible even after formaldehyde crosslinking of proteins. Initially, we tried to purify crosslinked chromatin by silica columns, however, the yield was almost zero (data not shown); therefore, we used silica magnetic beads instead of columns. **SPACE** - which stands for Silica Particle Assisted Chromatin Enrichment - **exploits the capacity of silica magnetic beads to purify formaldehyde-crosslinked chromatin in the presence of chaotropic salts** (Figure 1A). We prepared non-crosslinked negative controls in a similar way to routine DNA purification, which is normally free of contaminating proteins. We ran the proteins in the lysis buffer, washing buffers, the non-crosslinked control, and the crosslinked sample on an SDS-PAGE to check the stringency of the washes (Supplementary Figure 1A). By applying SILAC-labelling and mass spectrometry, crosslinked samples and non-crosslinked controls are pooled before adding silica magnetic beads. Thus, we are able to determine whether a protein is isolated due to the crosslinking or non-specific associations to the beads and other proteins.

**Figure 1:**
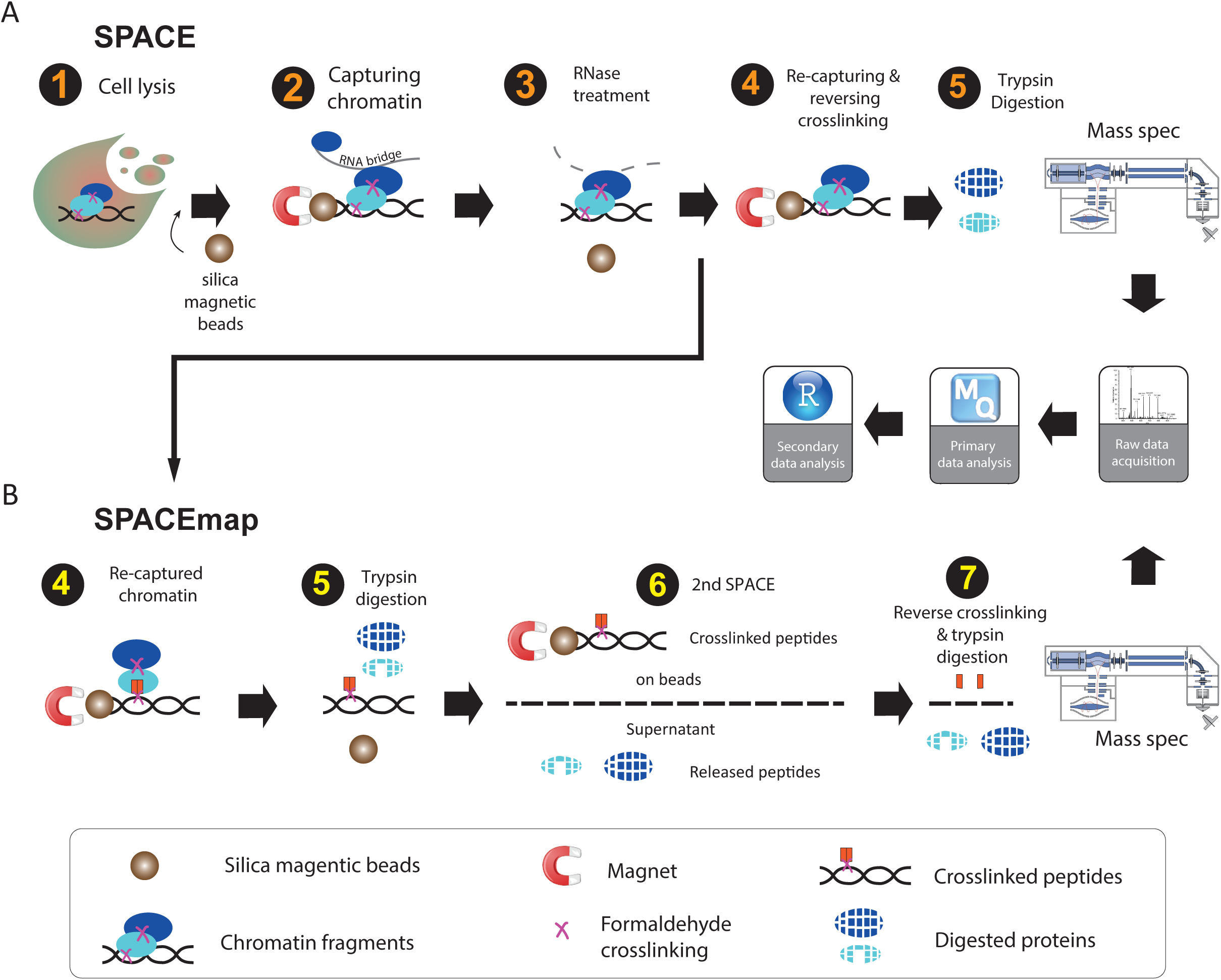
Overview of SPACE and SPACEmap. (A) In SPACE, 1: Cells are crosslinked by 1% formaldehyde, and resuspended in the lysis buffer containing guanidinium, and iso-propanol. Then silica magnetic beads are added to the lysate. 2: Chromatin binds to the magnetic beads and is separated from the lysate. The beads are washed with lysis buffer and ethanol. 4: Chromatin is eluted by sonication and is treated with RNase A. 4: Chromatin is captured again on the beads to be washed again with ethanol and Acetonitrile. Then the crosslinking is reversed, and trypsin/LysC are added to digest the chromatin-associated proteins on the beads. (B) In SPACEmap, chromatin is recaptured in step 4, however, the crosslinking is not reversed. 5: trypsin is added to digest the chromatin-associated proteins. 6: using another round of SPACE released peptides are separated from crosslinked peptides. Both crosslinked fractions and released fractions are injected to the mass spec to be compared quantitatively. After mass spectrometry and data acquisition, the raw files are analysed by MaxQuant to identify and quantify the proteins. Further statistical, domain and GO analysis are performed using R in RStudio.

**SPACE is stringent, yet fast and flexible, and requires little starting material.** Starting with as few as 20,000 cells, SPACE takes approximately 1h from the cell lysis to the start of protein digestion; it employs denaturing reagents to efficiently remove contaminants (4M guanidinium isothiocyanate, 2% Sarkosyl, 80% ethanol and 100% acetonitrile) and extensive RNase treatment (RNase A, 100ug, 30min at 37 C) to remove RNA-dependent interactors. The method is readily extended to identify chromatin-binding sites of proteins by a two-step digestion strategy (SPACEmap, Figure 1B). Additionally, SPACE can be combined with SICAP (Selective Isolation of Chromatin-Associated Proteins) (19) as a double purification and highly stringent variation of the method (Supplementary Figure 1B), or with ChIP to identify co-localized protein on chromatin (ChIP-SPACE) which is explained subsequently.

### SPACE shows increased specificity and sensitivity in comparison to other methods

We first applied SPACE to mouse embryonic stem (mES) cells cultured in 2iL using 2 forward replicates (heavy SILAC crosslinked), and 6 reverse replicates (light SILAC crosslinked). We considered proteins quantified with at least 1 forward experiment and 1 reverse experiment (≥ 2 experiments in total) for statistical analysis. We identified 1,459 significantly enriched proteins (1,349 proteins with log2FC > 1, adj. p-value < 0.01 in addition to 110 proteins with log2FC > 1 and adj. p-value < 0.1) compared with the non-crosslinked controls (Figure 2A-B, Table S1_SPACE). We assessed the correlation between all replicates (Figure 2B), which ranged from 0.46 to 0.91 (median R = 0.66). We then rigorously characterised the enriched proteins using keyword searching in gene ontology terms and protein information obtained from UniProtand DAVID databases (Figure 2C and Supplementary Figure 2A). We considered proteins that are related to chromatin functions or processes as potential true positive (PTP) which comprise 83% of the enriched proteins based on relative iBAQ values (as an estimation of protein abundances). Apart from those, proteins involved in translation, metabolic process, cell adhesion, protein folding, protein transport and miscellaneous proteins that are not present in the nucleus make up 3.3% of the enriched proteins. We considered these proteins as potential false positive (PFP) as they are not known to be involved in chromatin-related processes. Thus, using SPACE potential true positive biological processes are enriched 25-fold over potentially false positive terms. As examples, we identified 45 proteins that are involved in pluripotency or ES cell processes, including Oct4, Sox2 and Nanog as the core circuitry of pluripotency. In addition, we identified 11 proteins that are part of the polycomb group proteins (Supplementary Figure 2A).

**Figure 2:**
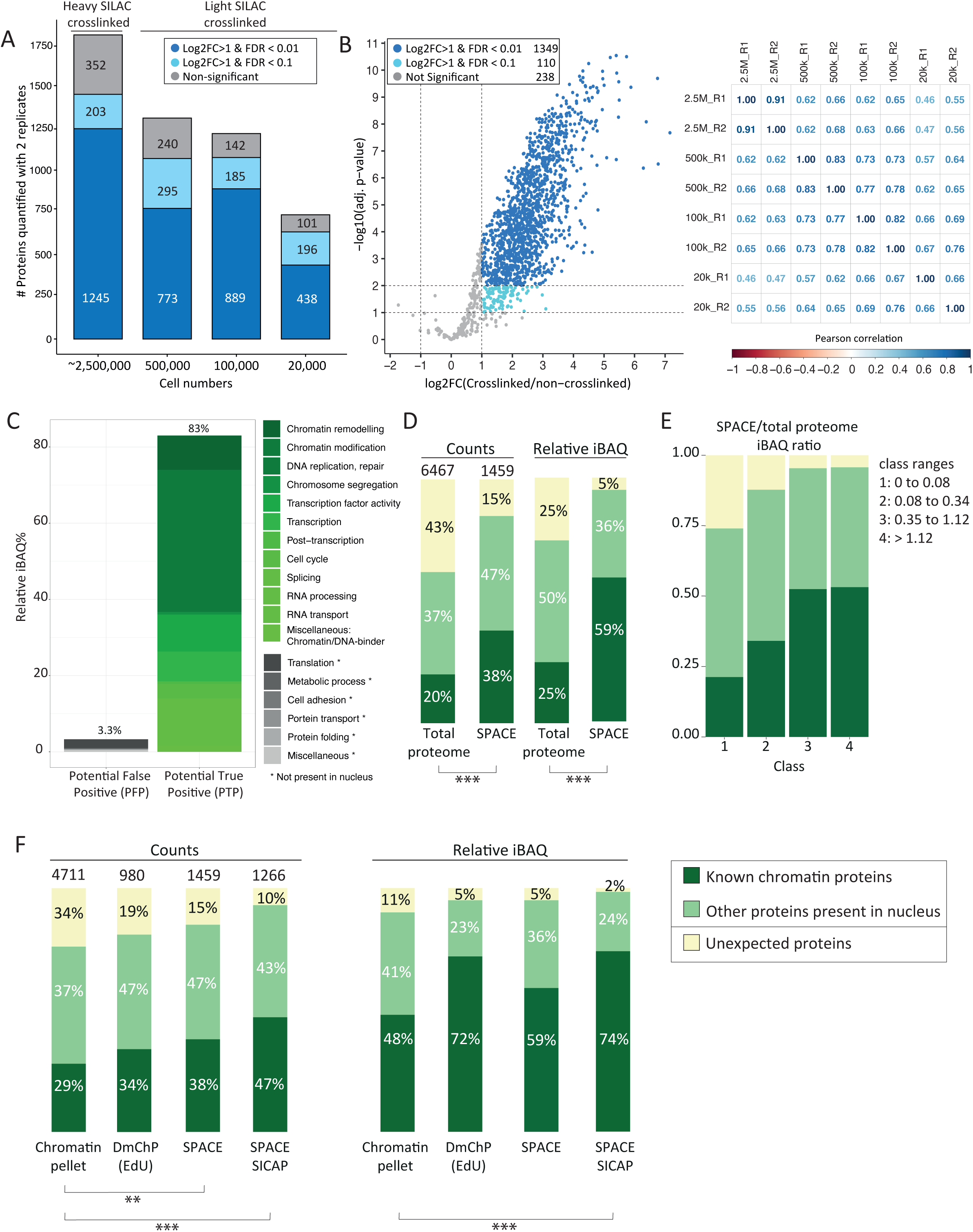
Chromatin composition in mES cells identified by SPACE. (A) SPACE experiments were carried out by varying number of cells. Each experiment was repeated twice. The bars show the proteins quantified by both replicates using each cell number. The dark blue stacks are very significantly enriched in comparison to the non-crosslinked control (adj. p-value < 0.01 and log2FC >1). The blue stacks are significantly enriched in comparison to the non-crosslinked control (adj. p-value < 0.1 and log2FC >1). The grey stacks are not significantly enriched. (B) All the experiments were integrated, and proteins quantified by at least 1 heavy SILAC crosslinked and 1 light SILAC crosslinked were considered for statistical analysis. The volcano plot shows the proteins that are very significantly enriched in comparison to the non-crosslinked controls with (adj. p-value < 0.01 and log2FC >1), proteins that are significantly enriched in comparison to the non-crosslinked controls (adj. p-value < 0.1 and log2FC >1) and proteins that are not significantly enriched with dark blue, blue, and grey, respectively. The matrix shows the Pearson correlations coefficient among the experiments. (C) The enriched proteins by SPACE were categorized based on their biological processes into potential true positive and potential false positive. (D) The enriched proteins were categorized into 3 groups: 1-‘known DNA or chromatin binding proteins’ (dark green), 2-‘Other proteins present in nucleus’ (pale green), 3-Proteins that do not fall into the previous categories are so-called ‘unexpected’ (yellow). The left two bars compare protein counts between the total proteome of mES cells and SPACE. The right two bars compare the relative iBAQ of the proteins. The total proteome data was obtained from published data (19), and re-analysed. Fisher’s exact test was used to show the statistical differences. (E) SPACE/total proteome iBAQ ratios for each protein was calculated. The proteins were classified into 4 equal groups based their ratios. The frequency of 1-‘known DNA/chromatin-binders’ (dark green), 2-‘proteins present in Nucleus’ (pale green) and 3-‘unexpected’ proteins (yellow), was shown in each class. (F) Chromatin pelleting, DmChP, SPACE and SPACE-SICAP results of mES cells were compared based on the protein counts and relative iBAQ of the enriched proteins. Chromatin pelleting (20) and DmChP (17) data were obtained from published data. The enriched proteins were categorized into 3 groups, as mentioned previously in panel B. Fisher’s exact test was used to show the statistical differences: *** p-value < 0.001, ** p-value ≤ 0.01 and * p-value ≤ 0.05

To evaluate the specificity of the method, we grouped proteins into three categories based on their gene ontology annotations (Figure 2D): 1) 554 (38%) known DNA or chromatin-binding proteins; 2) 686 (47%) proteins present in the nucleus but not annotated as DNA- or chromatin-binders; 3) and 219 (15%) other “unexpected” proteins, a large proportion of which are involved in translation. Weighted by relative iBAQ, it is apparent that known chromatin-binding proteins and proteins present in the nucleus are most abundant in the enriched proteins (59% and 36% respectively; Figure 2D), and the unexpected proteins have relatively low abundances (5%). Compared with the 6,467 proteins detected in the total proteome of whole-cell lysates (total proteome), **SPACE clearly enriches for canonical chromatin proteins, with additional representation of nuclear proteins that have not been previously identified to bind chromatin**.

Moreover, we compared the proteins that are compositionally biased due to the basic aminoacid or IDRs in their structure (reference = UniProt) between total proteome and SPACE (Supplementary Figure 2B). As a result, SPACE proteins are more enriched in basic aminoacids and IDRs in comparison to the total proteome.

We also calculated SPACE/total proteome iBAQ ratios to estimate how abundantly a given protein binds to chromatin (Figure 2E). We classified the proteins into 4 groups, with the higher SPACE/total proteome ratio the higher class. Interestingly, class 3 and 4 are more enriched in known DNA-chromatin-binding proteins, and they contain less unexpected proteins. In other words, having a high SPACE/total proteome ratio for a given protein suggests a high chromatin-binding chance. Nevertheless, relatively low SPACE/total proteome ratios should not be considered as a disproving of chromatin-binding ‘per se’. It is possible that a protein of interest is not efficiently crosslinked to chromatin, and it is partially removed during the purification procedure.

**To be more rigorous, we established an extremely stringent SPACE-SICAP double purification strategy:** the initial SPACE purification is followed by SICAP in which DNA is biotinylated with terminal deoxynucleotidyl transferase and captured by protease-resistant streptavidin magnetic beads (Supplementary Figure 1). SPACE-SICAP enriched 1,266 enriched proteins by at least 2 replicates, about ∼13% less than SPACE alone (Supplementary Figure 2C-D and Table S1_SPACE-SICAP). A DNase-treated control confirmed that the identification of chromatin-associated proteins depends on the presence of DNA: just 138 proteins were found, of which 101 were RBPs (Supplementary Figure 2D). We identified 908 proteins as the intersect of SPACE and SPACE-SICAP proteins (Supplementary Figure 2E).

The traditional method for chromatin isolation is subcellular fraction and centrifuge-assisted chromatin sedimentation. A recent method established to obtain a global view of chromatin composition is DmChP, which is based on prolonged EdU labelling to pull down DNA using Click chemistry. We compared chromatin pelleting (20), DmChP (17), SPACE and SPACE-SICAP to evaluate their specificity and sensitivity for isolating chromatin proteins from mES cells. As described previously, the proteins were categorized to 1) known DNA or chromatin-binders; 2) proteins known to be present in the nucleus but not annotated as DNA- or chromatin-binders; and 3) “unexpected” proteins. In addition to the number of the proteins, it is important to consider the abundance of the proteins to have a comprehensive view of specificity. We made the comparison based on protein counts and relative iBAQ values (as an estimation of protein abundance), and we used Fisher’s exact test to show the significant statistical differences. Based on protein counts, SPACE and SPACE-SICAP have better performance in isolating relevant proteins and removing unexpected proteins in comparison to chromatin pelleting, as evident by Fisher’s exact test (Figure 2F).

Statistically, we didn’t observe significant differences in the specificity of SPACE, SPACE-SICAP and DmChP (Figure 2F). While number of enriched proteins using SPACE is ∼50% more than DmChP (1,459 versus 982 enriched proteins), input material for SPACE is >10-fold less than DmChP (30 million versus 2.5 million cells per replicate for DmChP and SPACE, respectively). This indicates SPACE is more sensitive for chromatome studies which is not surprising, because SPACE doesn’t necessitate EdU labelling of DNA, Click chemistry and streptavidin pull down.

Limitation of input material is a burden for many chromatin proteomic studies, especially those using primary tissue samples or cell sorting. We, therefore, aimed to assess the sensitivity of SPACE by progressively decreasing the number of input cells from ∼2,500,000, 500,000, 100,000 and finally 20,000. We identified a reduced, but still substantial, number of proteins. The distribution of enriched proteins between ‘known chromatin proteins’, ‘present in the nucleus’ and ‘unexpected’ categories are very similar among these samples (Figure 2A and Supplementary Figure 2F). **Thus, SPACE is accurate and sensitive enough to be used for chromatome studies with limited input material**.

### SPACE reveals RBPs as major chromatin components

Strikingly, RBPs comprise a large proportion of the enriched proteins. Based on GO molecular functions, 696 RBPs are found in the SPACE dataset (48% of the enriched proteins), which comprise 74% of the enriched proteins weighted by iBAQ (Figure 3A). In other words, our SPACE data reveals 487 new caRBPs in addition to 209 previously characterised caRBPs. To understand if the RBPs in our dataset are associated with newly transcribed RNAs, we compared our results with RICK (23) and CARIC results (24) (Figure 3B and Supplementary Figure 3A). Both of these methods work by incorporating Ethynyl Uridine (EU) into the newly synthesized RNA. Then UV-crosslinking is applied to crosslink the RBPs to RNA, and nascent RNAs are captured using Click-chemistry. Interestingly, ∼43% (244+272+118 = 634, Supplementary Table S1) of the enriched proteins by SPACE data overlap with RICK and CARIC. However, some of these proteins are not annotated as RBPs based on GO molecular functions.

**Figure 3:**
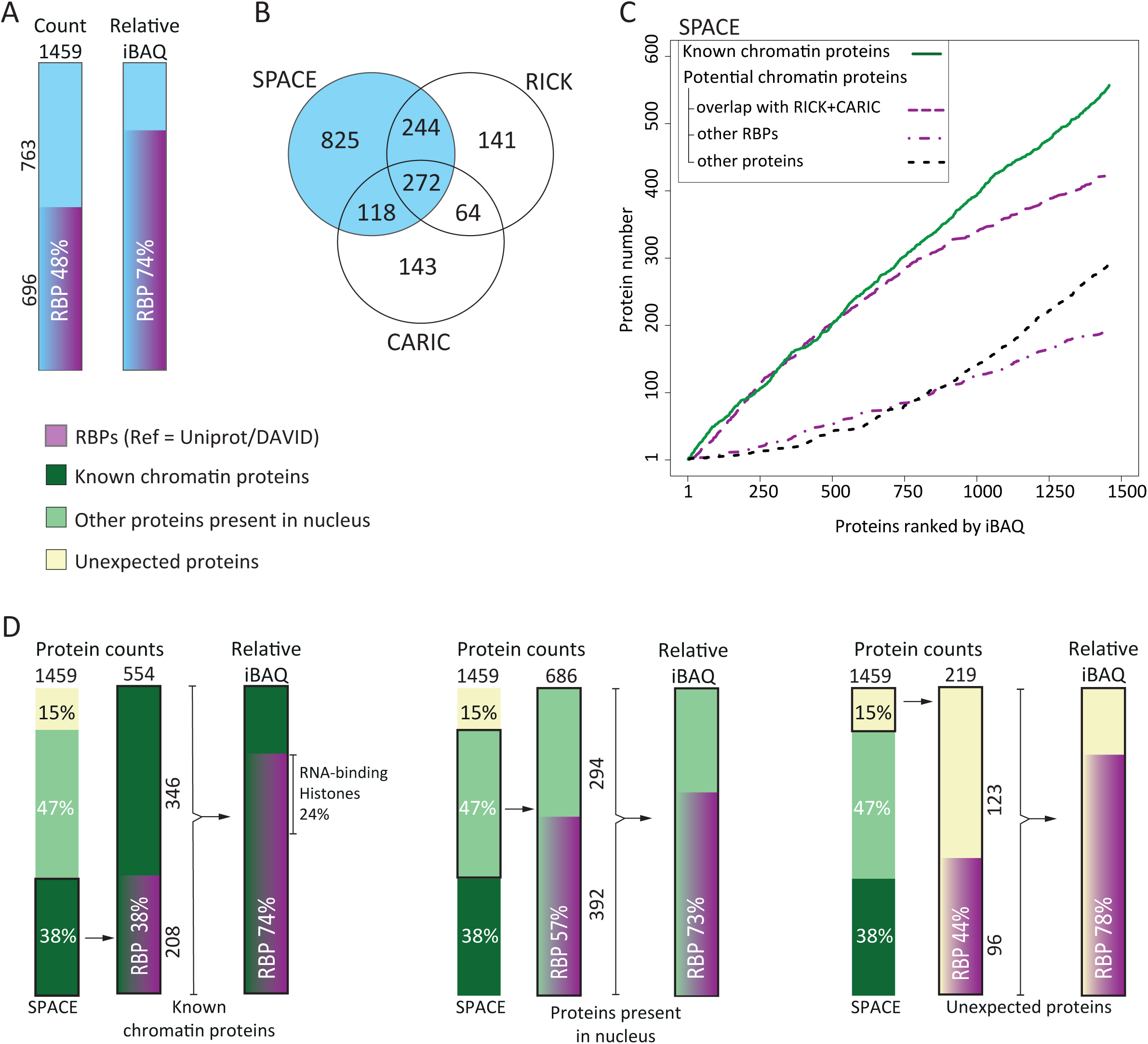
SPACE reveals RBPs as a major component of chromatome. (A) The frequency of RBPs in the entire proteins enriched by SPACE were shown as counts and relative iBAQ. (B) Comparing proteins enriched by SPACE with RICK(23) and CARIC(24) datasets. The latter two datasets enrich RBPs that interact with newly transcribed RNA. (C) Enriched proteins by SPACE were ranked by their relative iBAQ. The rates of accumulation in the dataset were compared among 1-known DNA/Chromatin-binders, including known caRBPs (continuous green line), 2-proteins that overlap with RICK and CARIC (dash purple line), 3-other RBPs (dot-dash purple line), and 4-other proteins (dot black line). (D) The proportion of RBPs in DNA/chromatin-binding proteins (left bars), protein present in nucleus (middle bars), and unexpected proteins (right bars) were shown as count and relative iBAQ.

To compare the estimated abundance of nascent-RBPs with the other enriched proteins, we then ranked the SPACE proteins based on their iBAQ values and compared the accumulation of 1-known chromatin proteins (including known chromatin-binder RBPs), 2-the overlapping proteins with RICK and CARIC, 3-other RBPs and 4-other proteins (Figure 3C). **Interestingly, proteins in group 2 (proteins associated with newly synthesized RNAs) are ranked higher than group 3 and group 4; indicating that they are more abundant in the context of chromatin**.

Among the known chromatin-binders in SPACE proteins, RBPs comprise 38% of the protein count (208 out of 554) and 74% by relative iBAQ. Focusing on the “proteins present in nucleus”, we find that RBPs comprise 57% and 73% by counts and abundance, respectively. Finally, there are 96 RBPs among 219 “unexpected” proteins which comprise 78% by relative iBAQ (Figure 3D).

We also developed SPACE-SICAP as a more stringent version of SPACE. We considered 908 proteins which are common to SPACE and SPACE-SICAP datasets (Supplementary Figure 3 B-C). **Again, we observed a strong enrichment of RBPs among chromatin-associated proteins, as 53% of the 908 proteins are RBPs. Altogether this result indicates dual DNA- and RNA-binding functionality in chromatin-associated proteins**. To inspect RBP interactions with chromatin more thoroughly we sought to identify chromatin-binding sites of RBPs.

### SPACEmap locates the specific chromatin-binding regions of proteins

To better understand how proteins are integrated into chromatin, we took an approach similar to RBDmap that identifies peptides crosslinked to RNA (25). However, instead of digesting the proteins with LysC or ArgC and then trypsin, we treated them twice with trypsin. Trypsin cleaves at both argininyl and lysinyl residues, so more peptides are digested and released in the first step, allowing us to identify crosslinked sites at higher resolution. Further, we used formaldehyde crosslinking, which is reversible (instead of UV-crosslinking used in RBDmap) which allowed for straightforward mass spec analysis.

To separate peptides crosslinked to DNA (crosslinked fraction), we digest proteins using large amounts of trypsin without reversing the crosslinking. As a result, most of the proteins are degraded and their peptides are released from the proteins (released fraction). Thus, crosslinked parts of the proteins to chromatin are purified (Figure 1B). We then carried out another round of SPACE, we heated the samples to reverse the crosslinking, and to detach the peptides from DNA in the crosslinked fraction. Both fractions were digested again by trypsin and compared with each other to identify the peptides that were significantly enriched in each fraction. Peptides enriched in the crosslinked fraction are either crosslinked directly to DNA, or indirectly via another peptide to DNA (Supplementary Figure 4A). Peptides indirectly crosslinked to DNA remain in the crosslinked fraction if the bridging peptides are long enough to connect DNA to the other peptides. In addition, 2 crosslinking sites are needed to build the bridge. Therefore, we anticipate the chance of enriching indirectly crosslinked peptides to DNA is lower than directly crosslinked peptides to DNA. In both cases, the peptides enriched in the crosslinked fraction are considered as the contact sites of the proteins with chromatin.

We identified 20,896 peptides, of which 6,158 were enriched in the crosslinked fraction and 6,312 in the released fraction (adj. p-value < 0.1 and log2FC > 0.4, Figure 4A). 4,420 peptides from 1,186 proteins were captured by the original SPACE method and in the crosslinked fraction of SPACEmap (Figure 4B, Table S2_SPACEmap peptides). Of these, ∼90% (3,956 peptides) mapped to a known protein domain or predicted intrinsically disordered region (IDR) or both.

**Figure 4:**
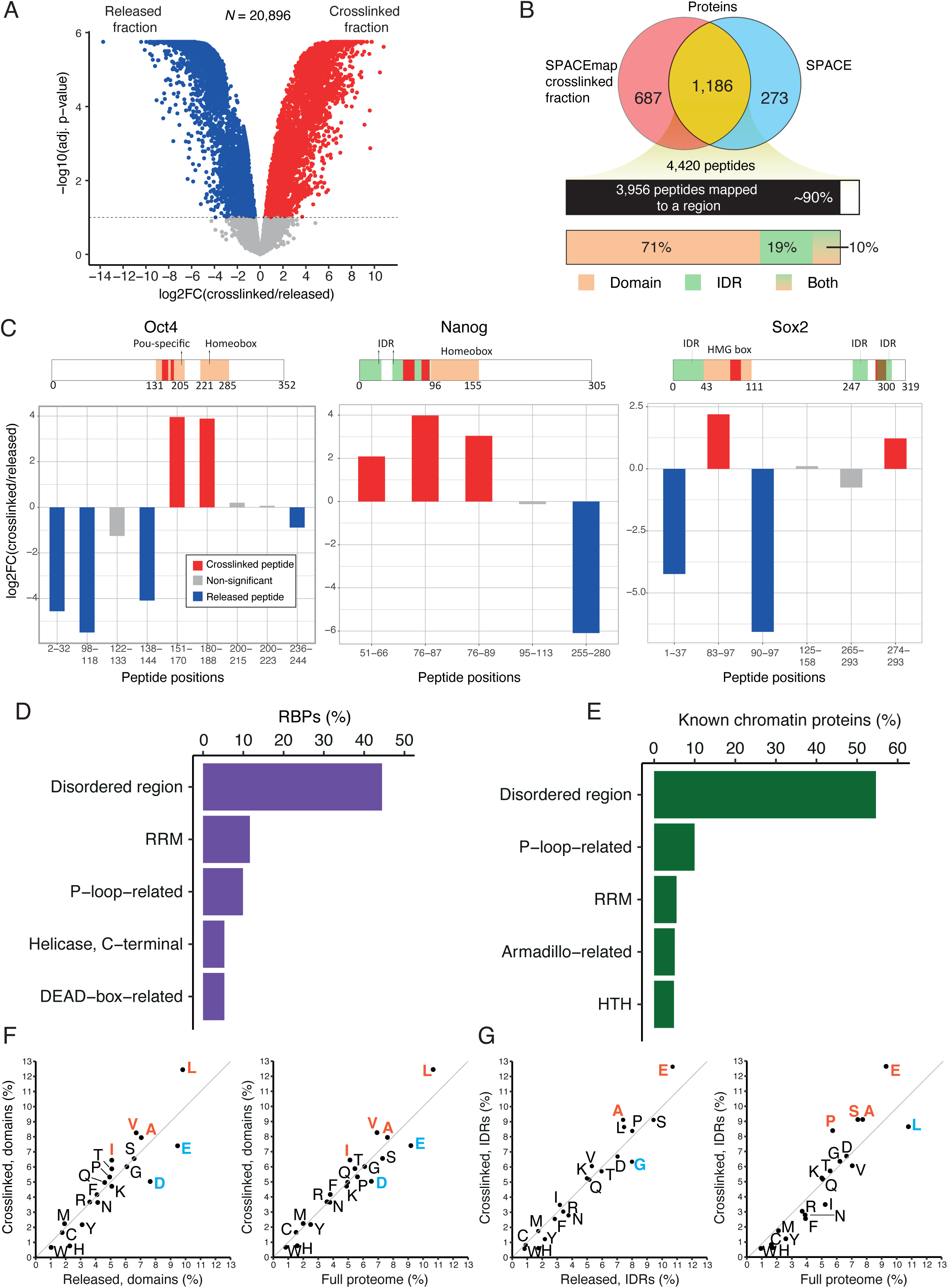
Locating chromatin-binding sites of the proteins. (A) The volcano-plot shows peptides enriched (adj. p-value < 0.1 and log2FC > 0.4) in the crosslinked fraction (red) and in the released fraction (dark blue). (B) The overlap of the proteins identified by crosslinked fraction (red) and SPACE (light blue) is shown in yellow. The upper bar shows the number of peptides corresponding to the overlapping proteins and the proportion of the peptides that are mapped to any regions (domains or IDRs). The lower bar shows the proportions of peptides that mapped to a domain, an IDR or both. (C) The plots show crosslinked and released peptides in Oct4, Sox2 and Nanog. The peptides significantly enriched in the crosslinked and released fractions are red and blue, respectively. Non-significantly enriched peptides are grey. The pink bars indicate the aminoacid positions of the DNA-binding domains. The green bars denote IDRs. The red boxes show the enriched peptides in the crosslinked fraction. (D) Top 5 domains/regions by the proportion of RBPs that contain them. RBPs containing these domains/regions have peptides enriched in the crosslinked fraction and overlapping with these domains/regions or residing no farther than 10 amino acids from them. (E) Top 5 domains/regions by the proportion of known DNA/chromatin proteins that contain them. The proteins containing these domains/regions have peptides enriched in the crosslinked fraction and overlapping with these domains/regions or residing no farther than 10 amino acids from them. (F) Aminoacid composition of the peptides mapped to domains in the crosslinked fraction relative to the peptides mapped to domains in the released fraction (left) and total proteome (right). These peptides overlap with domains or reside no farther than 10 amino acids from them. (G) Aminoacid composition of the peptides mapped to IDRs in the crosslinked fraction relative to the peptides mapped to domains in the released fraction (left) and total proteome (right). These peptides overlap with domains or reside no farther than 10 amino acids from them.

We compared the peptides from Oct4 (Pou5f1), Sox2 and Nanog with annotations of their DNA-binding regions (Figure 4C). The POU-specific domain of Oct4 extends from residues 131-205 (UniProt coordinates), and the precise DNA-binding residues are at positions 150, 157, 173-179 and 186-189 (Esch et al., 2013). Two peptides corresponding to positions 151-170 and 180-188 containing almost all the DNA-binding residues are enriched in the cross-linked fraction. Seven other peptides from the non-DNA-binding regions of Oct4 were not enriched. **Thus, the Oct4 peptides in the crosslinked fraction accurately match with Oct4’s known DNA-binding sites** (Figure 4C, left).

Nanog harbours a Homeobox domain that extends from residues 96 to 155. We identified three enriched peptides corresponding to positions 51-66, 76-87 and 76-89 (Figure 4C, middle). **All three peptides are located in the IDR adjacent to the homeodomain at the N-terminal region of Nanog (**Supplementary Figure 4B). The crystal structure of the Nanog homeodomain suggests protein-DNA interface is located between residues 136-152-Helix H3 (26); here, we lack tryptic peptides encompassing this region owing to the large number of lysine and arginine residues. Our result suggests there is a protein-chromatin interface in the IDR close to the homeodomain. Thus, whereas crystal structures provide detailed information about interactions involving ordered protein regions, SPACEmap complements with insights into chromatin interactions from IDRs which might otherwise be missed.

Finally, Sox2 contains an HMG box domain located at residues 43-111. We identified six Sox2 peptides, two of which were enriched in the crosslinked fraction. **The peptide encompassing residues 83-97 is located within the HMG box, whereas the peptide from residues 274-293 is located in the IDR of Sox2 near the C-terminus of the protein** (Figure 4C, right, and Supplementary Figure 4C). Our result predicts an additional chromatin-interacting element near the C-terminal domain of Sox2 (274-293).

Subsequently, we examined crosslinked fraction at peptide and protein levels to understand how RBPs bind to chromatin. **We found that ∼44% of RBPs have at least one crosslinked peptide that maps to IDRs** (Figure 4D, Supplementary Figure 4D, Table S2_peptides mapped to a region). Strikingly, ∼55% of ‘known chromatin proteins’ have at least one crosslinked peptide that maps to IDRs (Figure 4E, Supplementary Figure 4E). A recent study (27) has indicated that IDRs interact with DNA using low-affinity interactions also interfacing with histones. Initially, IDR-guided weak interactions may allow accelerated recognition of broad DNA regions. Subsequently, DNA-binding domains could stably bind to specific DNA motifs (27).

We also observed p-loop domains among the top 5 enriched domains (Figure 4D-E). Although p-loop domains are associated with phosphate-binding such as nucleotide-triphosphates (NTPs), they emerged as avid RNA-binding and ssDNA-binding domains (28). As such, **our result confirms p-loop interactions with chromatin in living cells**. In addition, classical RNA or DNA-binding domains such as RRM, helicases and helix-turn-helix (HTH) domains are highly enriched in the crosslinked fraction (Figure 4D-E).

To further understand the general characteristics of crosslinked fraction peptides, we compared their amino acid composition with the released fraction peptides, as well as the peptides from the total proteome. Negatively charged residues glutamate and aspartate are depleted in the crosslinked fraction peptides that map to the domains, whereas **hydrophobic residues such as leucine, valine, alanine and isoleucine are enriched** (Figure 4F). **The crosslinked fraction peptides that map to IDRs are enriched in glutamate, as well as proline** (Figure 4G), which agrees with the fact that proline and glutamate are the most disorder-promoting residues (29). It is surprising that glutamate is depleted from crosslinked peptides mapped to domains but enriched in those mapped to the IDRs. It is likely that the glutamate residues in the IDRs are involved in protein-protein interactions on chromatin. Alternatively, glutamate residues may destabilize the interactions between the proteins and the target binding sites on DNA to accelerate target recognition. Yet, the precise role of glutamate or proline in interactions between IDRs and DNA or chromatin remains to be understood.

During the SPACEmap procedure, the crosslinked protein complexes are broken down, and only peptides remain crosslinked to DNA. As a result, abundant proteins are removed more efficiently, and they are prohibited from associating with DNA during the purification procedure. Therefore, SPACEmap is even more stringent than SPACE for identification of chromatin-binding proteins. Intersecting SPACE and the crosslinked fraction hits yielded 1,186 proteins (Figure 4B). Among them, we found 598 RBPs of which 194 proteins were previously known as DNA/chromatin-binders. Thus, SPACEmap provides strong evidence of chromatin-binding for 404 RBPs (Table S2_SPACEmap-verified caRBPs). **Altogether, SPACEmap stringently verifies chromatin-binding proteins and faithfully detects their chromatin interface**.

### SPACE elucidates features of mES cells in the ground and metastable states

To demonstrate the quantitative capacity of SPACE, we compared mES cells grown in 2iL (the ground-state) and serum medium (the metastable state) in order to identify caRBPs in different pluripotency conditions. We identified 1,880 proteins in total (Figure 5A): 100 proteins were significantly more abundant in 2iL and 87 in serum (Log2FC > 1 and adj. p-value < 0.1, Table S3_comparative SPACE). We also compared the SPACE results with the total proteome from the total cell lysate. We found 1,768 proteins in the intersection of SPACE and total proteome, and there was a strong correlation in log2 fold-change values between them (Figure 5B and supplementary Figure 5A-B; R = 0.62). This indicates chromatin-binding is largely regulated at the protein expression level. However, there are proteins that are differentially regulated at the level of chromatin-binding, while their expression (total amount) does not change (Figure 5B, the yellow lane). As an example, b-Catenin binds to chromatin in 2iL medium ∼ 3-fold higher than serum condition. While, in total b-Catenin is up-regulated ∼1.5-fold. Thus, activation of Wnt pathway by inhibiting Gsk3b (CHIR99021) is significantly detectable by SPACE.

**Figure 5:**
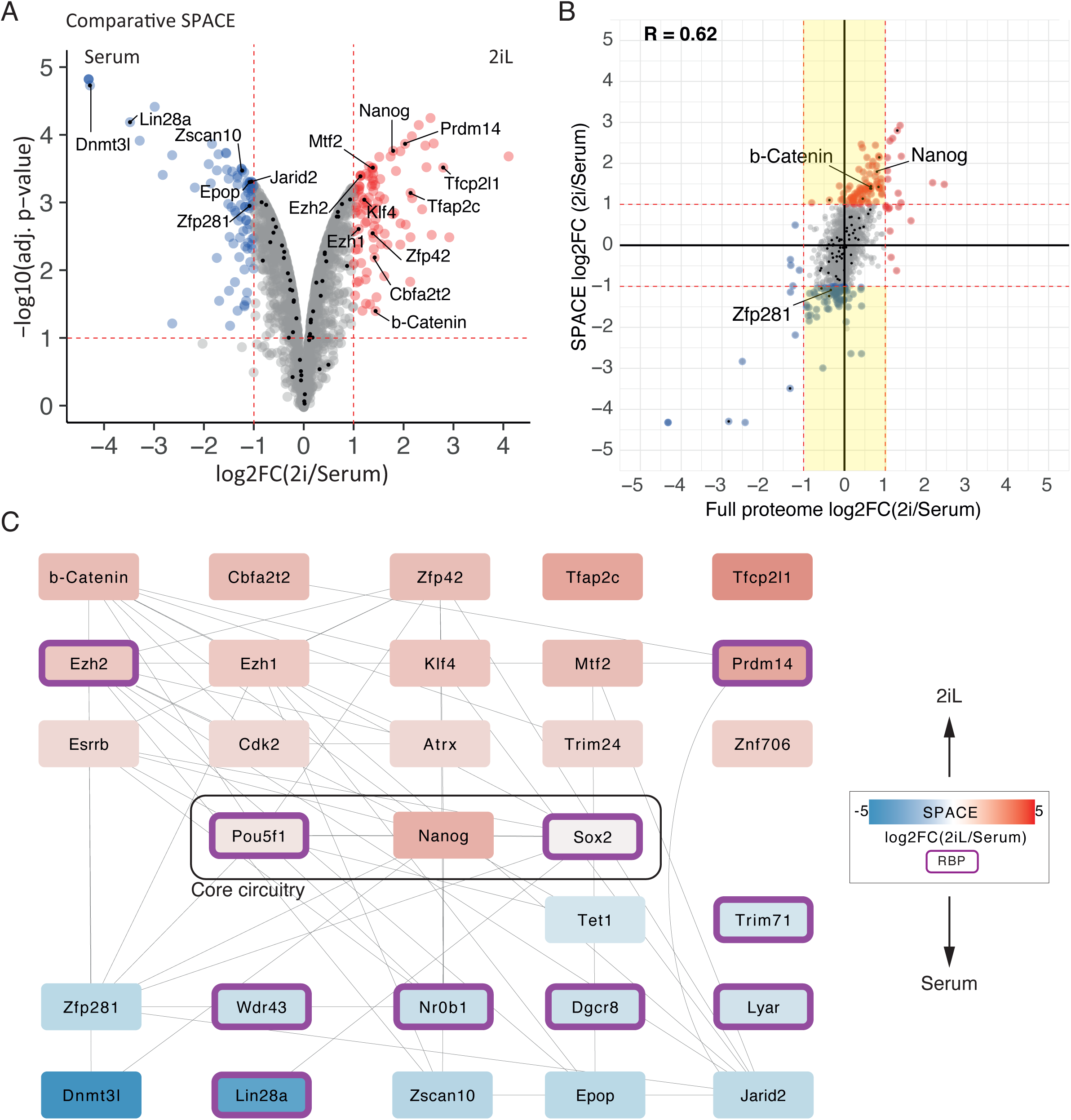
Chromatin composition in 2iL and serum conditions of mES cells. (A) The volcano-plot shows proteins that are significantly more abundant in 2iL and serum by red and blue, respectively (adj. p-value <0.1 and log2FC >1). The rest of the proteins were depicted by grey. Proteins involved in pluripotency, mES cell self-renewal or differentiation were marked by black dots. (B) Comparing total proteome analysis with SPACE. The yellow lane indicates differentially regulated proteins detectable only by SPACE. The total proteome data was obtained from (19), and re-analysed. (C) Experimental interaction network of the proteins involved in pluripotency, mES cell self-renewal or differentiation. RBPs were marked in purple borders.

To understand how the global network of pluripotency is regulated in 2iL and serum conditions, we looked for proteins with known functions in maintaining embryonic stem cells or exiting from pluripotency. We identified 68 proteins that are positively or negatively involved in the self-renewal of pluripotent stem cells. The network in Figure 5C depicts previously known experimental interactions between a subset of them (Log2FC > 0.6 and adj. p-value < 0.1). Among them are chromatin proteins that physically interact with the core circuitry of pluripotency (Nanog, Oct4, Sox2). Our data suggests that the network of protein interactions surrounding the core pluripotency circuitry shifts substantially between the 2iL and serum conditions. In agreement with previous studies, our results indicate that Tfcp2l1, Prdm14, Cbfa2t2, Zfp42 (Rex1), Klf4, Trim24 and Esrrb (30,31) bind to chromatin preferentially in 2iL conditions, whereas Lin28a and Zfp281 bind more abundantly to chromatin in serum conditions. Our results are in line with the role of Lin28a and Zfp281 in transitioning from naive to primed state of pluripotency (32,33). Interestingly, differential regulation of Zfp281 is only detectable by SPACE but not total proteome (Figure 5B). Thus, SPACE reveals how the ES cells respond to the cellular conditions more thoroughly than a total proteome analysis. The reason is that SPACE measures both quantity of the proteins, and their binding to chromatin. While a total proteome analysis measures only the quantity of the proteins.

Among the differentially enriched proteins there are 70 RBPs (adj. p-value < 0.1 and log2FC > 1, Supplementary Figure 5C). Lin28a is a well-characterised RBP that prevents ES cell differentiation by suppressing let-7 (34). Together with Prdm14, they are also known for their roles in DNA-demethylation by recruiting Tet proteins in mouse ES cells; thus, their presence among chromatin-binders was expected (35,36). Our data also indicates Dazl as a caRBPs with highly differential chromatin-binding ability (log2FC > 2) in 2iL condition. Additionally, Dazl has a very high SPACE/total proteome iBAQ ratio (1.55, Supplementary Figure 5D). These findings led us to examine Dazl’s chromatin-binding by other methods.

### Dazl a 3’-UTR-binding protein is recruited to transcription start sites on chromatin

Dazl is best known for targeting the 3’ untranslated regions (3’ UTRs) of mRNAs to regulate their translation, especially in germ cells (37,38). We first assessed Dazl’s cellular localization by immunofluorescent staining using a validated antibody, which confirmed that it is present both in the nuclei and cytoplasm of mES cells (Supplementary Figure 6A). We then performed chromatin immunoprecipitation and sequencing (ChIP-seq) to investigate the genome-wide locations of Dazl binding sites Figure 6A), revealing ∼1,300 reproducible peaks. Considering Dazl has known 3’ UTR-binding properties, we were surprised to find that 75% of peaks are found within a 1kb of window centred on transcription start sites (TSS); many target genes are developmental regulators, including Hox genes (Supplementary Figure 6B), several Wnt ligands and Frizzled receptors. As most of the Dazl target genes are involved in development and differentiation of mES cells, we compared Dazl, Suz12, Aebp2 and H3K27me3 profiles (Figure 6A the heatmap). Interestingly, we observed very similar binding patterns, demonstrating that **Dazl co-localizes with PRC2 on chromatin, especially at the promoters of genes related to the differentiation programs and exiting from pluripotency**.

**Figure 6:**
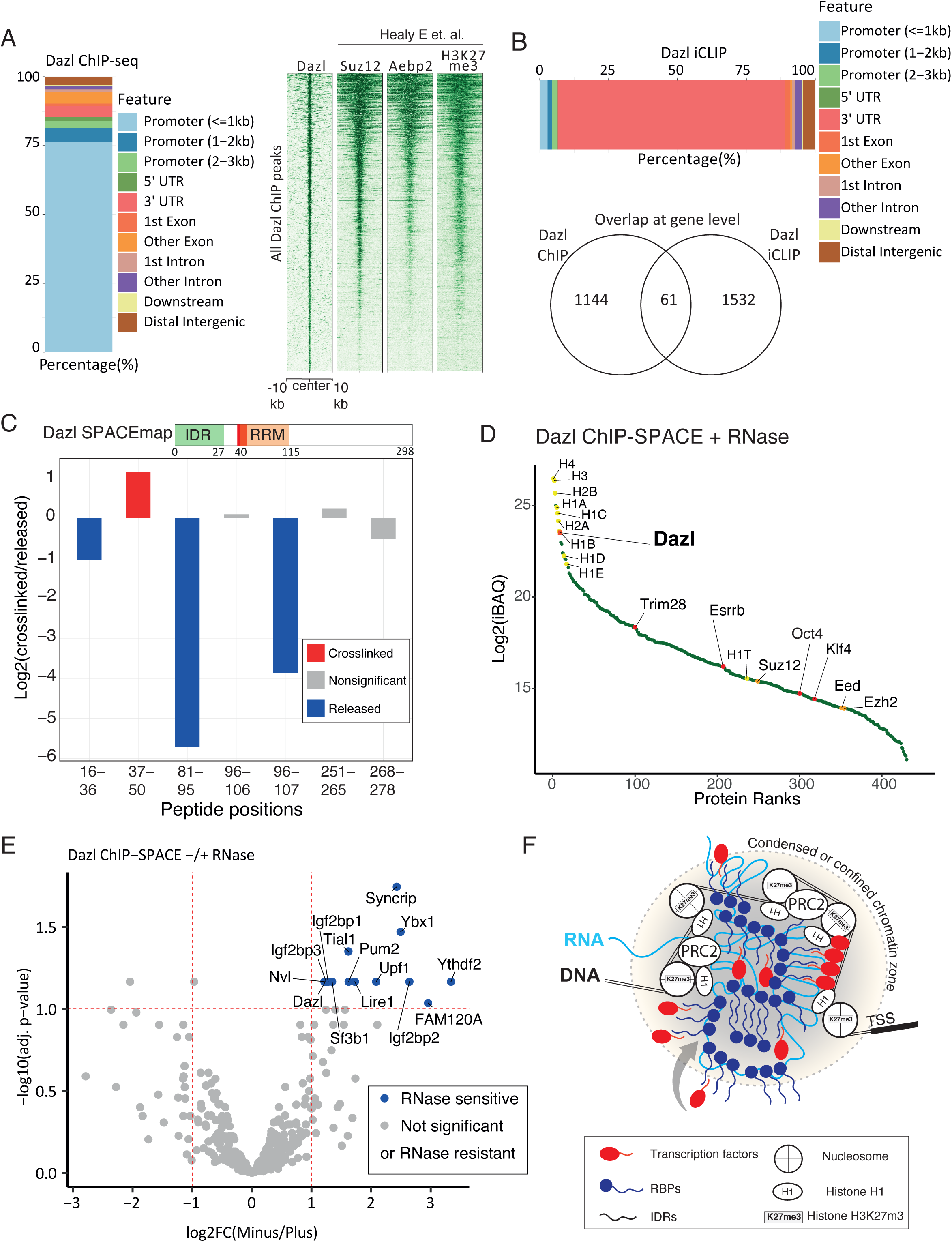
Dazl is recruited to silenced transcription start sites in mES cells. (A) Annotation of Dazl ChIP-seq peaks, and the profile of Dazl peaks on the genome in comparison with Suz12, Aebp2 and H3K27me3 peaks in mES cells. The last ChIP profiles of Suz12, Aebp2 and H3K27me3 were obtained from (21). (B) Annotation of Dazl iCLIP peaks (top bar), and the intersect of Dazl ChIP-seq and iCLIP-seq peaks at the gene level (bottom Venn diagram). (C) Dazl peptides identified using SPACEmap procedure are shown by red (enriched in crosslinked fraction), blue (enriched in released fraction) and grey (statistically non-significant). (D) Proteins enriched by Dazl ChIP-SPACE in comparison to the IgG control were sorted by the abundance of the proteins (iBAQ). Histones and PRC2 components are shown by yellow and orange dots, respectively. Dazl and transcription factors are shown by red dots. (E) The volcano plot shows proteins identified by ChIP-SPACE and their sensitivity to RNase A treatment. Proteins that are affected by RNase treatment are named in the plot. (F) The schematic model of RBP interactions with chromatin based on Dazl data. Chromatin-associated RBPs form a condensed or confined zone probably via interactions among their IDRs with other components of chromatins. The RBPs that are in the periphery of the zone are sensitive to RNase treatment. The RBPs in the centre of the zone are resistant to RNase treatment. Transcription factors and other components of chromatin are probably recruited or trapped by the RBPs.

We also performed individual-nucleotide crosslinking and immunoprecipitation (iCLIP) to identify the RNA-binding sites of Dazl across the transcriptome (39). We identified 2,550 peaks in mRNAs, 2099 of which were found in 3’ UTRs, and only 166 located within 3,000 nucleotides of the 5’ end of mRNAs (Supplementary Table S4_Dazl ChIP, iCLIP and ChIP-SPACE). Thus, the RNA binding sites were positioned at different locations in genes compared to DNA-binding sites, which were located mainly in promoters (Figure 6B). Moreover, most of the genes containing DNA-binding sites of Dazl in their promoter or gene body did not overlap with the genes containing RNA-binding sites of Dazl within their transcripts; only 61 out of 1144 genes (5%) with a gene-proximal ChIP-seq-defined peak on their genes (gene body and 3kb upstream of the TSS) also have an iCLIP-defined peak on their respective transcripts. These results suggest that the **chromatin- and RNA-binding functions of Dazl are mechanistically independent**.

Next, we examined our SPACEmap data to understand how Dazl binds to chromatin. We observed that out of the 7 peptides that were present in SPACEmap data, only **one peptide was enriched in the crosslinked fraction, corresponding to Dazl’s RRM domain** (Figure 6C). RRM domains are known to participate in RNA-binding and DNA-binding; therefore, it remains to be seen whether Dazl binds to chromatin via a bridging RNA, or if it directly binds to the DNA itself. The first option might be plausible, despite the harsh RNase treatment, if RNA is incorporated into a multi-protein Dazl-containing complex that can partly protect it from RNase.

To study Dazl complexes on chromatin, we took a regional approach to identify proteins co-localised on chromatin with Dazl. Here, we developed ChIP-SPACE (Supplementary Figure 6C), a faster and less laborious method than ChIP-SICAP (19,40) as it excludes DNA end-labelling and streptavidin purification and used it to identify Dazl chromatin partners. Following ChIP, we treated our samples with and without RNase A, then purified chromatin fragments by SPACE. 442 proteins were enriched in comparison with the IgG control (moderated t-test BH adj. p-value < 0.1 and log2FC > 1, Figure 6D-E). Sorting the enriched proteins based on their abundance (iBAQ) revealed histones followed by Dazl as the most abundant proteins. In addition, we identified several histones H1, as well as three members of the PRC2 complex: Ezh2, Eed and Suz12. Moreover, we identified pluripotency transcription factors such as Oct4, Klf4, Trim28, Esrrb and 155 SPACEmap-verified caRBPs (∼35% out of 442). These findings indicate that **Dazl is part of a conglomerate of caRBPs and transcription factors that are colocalizing with PRC2 and the linker Histone H1 in the vicinity of TSSs (Figure 6F)**.

## Discussion

Here, we present SPACE, a robust, sensitive, and accurate method for purifying chromatin-associated proteins by silica magnetic beads for proteomic analysis. **Strikingly, SPACE revealed that ∼48% of the chromatome are potentially able to interact with RNA**. To identify the specific protein regions that participate in contacts with chromatin, we developed SPACEmap, which showed that ∼44% of the potential RBPs bind to chromatin via their IDRs. Similarly, according to RBDmap nearly half of the RNA-binding sites map to the IDRs (25). Proteins enriched in IDRs are essential for many chromatin functions such as transcriptional regulation and RNA processing (41). IDRs are primary drivers of phase separation of proteins into biomolecular condensates (12,42), which are important in organizing the local chromatin structure (43,44). Also, the activation domains of transcription factors consist of IDRs which enable transcription factors to phase separate with Mediator co-activators (45). A recent study has shown that IDRs can generate confinement states for transcription factors to increase the local concentration of transcription factors thereby altering transcriptional output (46). **Our findings demonstrate that RBPs directly interact with chromatin components, largely via their IDRs. Probably, RBPs contribute to the condensed or confined chromatin zone formation using their IDRs to recruit or trap transcription factors and other chromatin components (Figure 6F)**.

We compared the global chromatin composition in 2iL and serum conditions of mES cells, and **we observed Dazl as one of the most differentially expressed caRBPs**, which is highly upregulated on chromatin in the 2iL condition. Dazl has been primarily studied in the context of germ cells due to its substantial roles in controlling the mRNA translation and stability; especially mRNA of genes that are necessary for germ cell survival (37,47). To identify Dazl binding sites on chromatin we used ChIP-seq, and we found that Dazl associates with the same chromatin sites as PRC2. Thus, in contrast to a recent study that has shown RBPs often interact with enhancers, promoters and transcriptionally active regions on chromatin (2), our result indicates Dazl mostly binds to the transcriptionally silenced genes in mES cells (e.g. developmental genes). SPACEmap data reassures Dazl chromatin-binding and reveals Dazl’s RRM domain as the chromatin contact site. Our ChIP-SPACE result also indicates >1/3 of the proteins co-localized with Dazl on chromatin are RBPs; providing a large number of IDRs to drive condensate formation. In addition, there are 5 Histone H1 in the dataset together with the core nucleosomes. It has been shown that the disordered histone H1 tail forms phase separated condensates and behaves like a liquid glue that clamps condensed clusters of nucleosomes together (48,49). **Thus, our results suggest caRBPs can generate condensed chromatin zones which are transcriptionally silent**. Recently, an “RNA-bridge” model was proposed for PRC2 that requires RNA for proper chromatin localization (50). Conceivably, caRBPs bind to RNA-bridges to promote phase-separated PRC condensates and chromatin compaction. The precise role of caRBPs in phase separation-mediated PRC condensation remains to be elucidated.

In addition to Dazl, we found Lire1 as a RBP which binds to chromatin preferentially in serum condition. Lire1 is a nucleicacid-binding chaperone that mobilizes LINE-1 elements in the genome, and its differential regulation in serum condition and primed state pluripotency is highly intriguing and warrants further investigation.

SPACE is broadly applicable due to its superior sensitivity, as 100,000 cells are sufficient to enrich >1,000 chromatin associated proteins in a single-shot injection into the mass spec. We believe SPACE will be particularly valuable for quantitative comparisons in timepoint studies, or for analyses of microdissected or sorted cell types. Past studies required much larger amounts of material (20,51), and they required the incorporation of modified nucletoides such as EdU or biotin-dUTP into DNA (17,52,53). Many cell types, such as mES cells, are particularly sensitive to modified nucleotides (54). In addition, incorporation of modified nucleotides to tissues such as patient samples is impossible or hardly doable. Additionally, DNA replication is necessary for global incorporation of EdU into the genome. As such, EdU-based methods are limited to the actively proliferating cells. SPACE overcomes all these limitations, while also being more straightforward and highly sensitive.

Formaldehyde is widely used in the field of chromatin studies. The small molecules of formaldehyde connect groups that are ∼2 A□ apart (reviewed in (55)), thus formaldehyde crosslinking allows for capturing interactions between DNA-protein and protein-protein on chromatin. To avoid over-crosslinking, we applied formaldehyde in the medium of the cells. Thus, aminoacids of the medium compete with formaldehyde molecules. Nevertheless, the possibility of multi-indirect chromatin-binders should be considered. We, therefore, developed SPACEmap to make sure the RBPs are not over-crosslinked to chromatin (explained in Supplementary Figure 4A). As such, SPACEmap verified chromatin-binding of 404 RBPs, and determined their chromatin-contact regions.

All in all, our study demonstrates the capacity of SPACE for quantitative analyses of chromatin composition across conditions, and the capacity of SPACEmap to identify the regions of proteins that contact chromatin. Due to the ease of its application, its high sensitivity and specificity, these methods hold a great potential for further applications that could unravel the dynamics of gene regulation and genome maintenance in development and diseases. Specifically, studying neurodegeneration using SPACE and its variants will shed light on the mechanism of the disease, and reveal novel therapeutic approaches.

## Supporting information

Supplementary Material with figures

## Data availability

The mass spectrometry proteomics data have been deposited to the ProteomeXchange Consortium via the PRIDE partner repository with the dataset identifier PXD023903. The accession numbers for the Dazl ChIP-seq and iCLIP reported in this paper are ArrayExpress: E-MTAB-9302 and E-MTAB-9332, respectively.

## Acknowledgement

We would like to thank Jeroen Krijgsveld for revising the manuscript. We appreciate Flora Lee for sharing the iCLIP protocol. We would like to thank Yongkai Tan for his helps with handling next generation sequencing samples and data.

## Author contribution

MR, JU and NML designed the research. MR performed the SPACE, SPACE-SICAP, ChIP-SPACE and ChIP-seq experiments. MR analysed all the proteomics data. SS analysed SPACEmap data. JAZ performed Dazl ICC and iCLIP experiment and analysed the data. SS2 and MR analysed the ChIP-seq data. KD performed the SPACE-PAGE experiment. MR, JU and NML wrote the manuscript with input from all authors.

## Reference

1. Bentley, D.L. (2014) Coupling mRNA processing with transcription in time and space. Nat Rev Genet, 15, 163–175.

2. Xiao, R., Chen, J.Y., Liang, Z., Luo, D., Chen, G., Lu, Z.J., Chen, Y., Zhou, B., Li, H., Du, X. et al. (2019) Pervasive Chromatin-RNA Binding Protein Interactions Enable RNA-Based Regulation of Transcription. Cell, 178, 107–121 e118.

3. Oktaba, K., Zhang, W., Lotz, T.S., Jun, D.J., Lemke, S.B., Ng, S.P., Esposito, E., Levine, M. and Hilgers, V. (2015) ELAV links paused Pol II to alternative polyadenylation in the Drosophila nervous system. Mol Cell, 57, 341–348.

4. Herzel, L., Ottoz, D.S.M., Alpert, T. and Neugebauer, K.M. (2017) Splicing and transcription touch base: co-transcriptional spliceosome assembly and function. Nat Rev Mol Cell Biol, 18, 637–650.

5. Bregman, A., Avraham-Kelbert, M., Barkai, O., Duek, L., Guterman, A. and Choder, M. (2011) Promoter elements regulate cytoplasmic mRNA decay. Cell, 147, 1473–1483.

6. Trcek, T., Larson, D.R., Moldon, A., Query, C.C. and Singer, R.H. (2011) Single-molecule mRNA decay measurements reveal promoter-regulated mRNA stability in yeast. Cell, 147, 1484–1497.

7. Zid, B.M. and O’Shea, E.K. (2014) Promoter sequences direct cytoplasmic localization and translation of mRNAs during starvation in yeast. Nature, 514, 117–121.

8. Henninger, J.E., Oksuz, O., Shrinivas, K., Sagi, I., LeRoy, G., Zheng, M.M., Andrews, J.O., Zamudio, A.V., Lazaris, C., Hannett, N.M. et al. (2021) RNA-Mediated Feedback Control of Transcriptional Condensates. Cell, 184, 207–225 e224.

9. Shrinivas, K., Sabari, B.R., Coffey, E.L., Klein, I.A., Boija, A., Zamudio, A.V., Schuijers, J., Hannett, N.M., Sharp, P.A., Young, R.A. et al. (2019) Enhancer Features that Drive Formation of Transcriptional Condensates. Mol Cell, 75, 549–561 e547.

10. Banani, S.F., Lee, H.O., Hyman, A.A. and Rosen, M.K. (2017) Biomolecular condensates: organizers of cellular biochemistry. Nat Rev Mol Cell Biol, 18, 285–298.

11. Shukla, S. and Parker, R. (2016) Hypo- and Hyper-Assembly Diseases of RNA-Protein Complexes. Trends Mol Med, 22, 615–628.

12. Tauber, D., Tauber, G. and Parker, R. (2020) Mechanisms and Regulation of RNA Condensation in RNP Granule Formation. Trends Biochem Sci, 45, 764–778.

13. Gebauer, F., Schwarzl, T., Valcarcel, J. and Hentze, M.W. (2021) RNA-binding proteins in human genetic disease. Nat Rev Genet, 22, 185–198.

14. Van Nostrand, E.L., Freese, P., Pratt, G.A., Wang, X., Wei, X., Xiao, R., Blue, S.M., Chen, J.Y., Cody, N.A.L., Dominguez, D. et al. (2020) A large-scale binding and functional map of human RNA-binding proteins. Nature, 583, 711–719.

15. Shiio, Y., Eisenman, R.N., Yi, E.C., Donohoe, S., Goodlett, D.R. and Aebersold, R. (2003) Quantitative proteomic analysis of chromatin-associated factors. J Am Soc Mass Spectrom, 14, 696–703.

16. Sirbu, B.M., Couch, F.B., Feigerle, J.T., Bhaskara, S., Hiebert, S.W. and Cortez, D. (2011) Analysis of protein dynamics at active, stalled, and collapsed replication forks. Genes Dev, 25, 1320–1327.

17. Aranda, S., Alcaine-Colet, A., Blanco, E., Borras, E., Caillot, C., Sabido, E. and Di Croce, L. (2019) Chromatin capture links the metabolic enzyme AHCY to stem cell proliferation. Sci Adv, 5, eaav2448.

18. Ritchie, M.E., Phipson, B., Wu, D., Hu, Y., Law, C.W., Shi, W. and Smyth, G.K. (2015) limma powers differential expression analyses for RNA-sequencing and microarray studies. Nucleic Acids Res, 43, e47.

19. Rafiee, M.R., Girardot, C., Sigismondo, G. and Krijgsveld, J. (2016) Expanding the Circuitry of Pluripotency by Selective Isolation of Chromatin-Associated Proteins. Mol Cell, 64, 624–635.

20. van Mierlo, G., Dirks, R.A.M., De Clerck, L., Brinkman, A.B., Huth, M., Kloet, S.L., Saksouk, N., Kroeze, L.I., Willems, S., Farlik, M. et al. (2019) Integrative Proteomic Profiling Reveals PRC2-Dependent Epigenetic Crosstalk Maintains Ground-State Pluripotency. Cell Stem Cell, 24, 123–137 e128.

21. Healy, E., Mucha, M., Glancy, E., Fitzpatrick, D.J., Conway, E., Neikes, H.K., Monger, C., Van Mierlo, G., Baltissen, M.P., Koseki, Y. et al. (2019) PRC2.1 and PRC2.2 Synergize to Coordinate H3K27 Trimethylation. Mol Cell, 76, 437–452 e436.

22. Blazquez, L., Emmett, W., Faraway, R., Pineda, J.M.B., Bajew, S., Gohr, A., Haberman, N., Sibley, C.R., Bradley, R.K., Irimia, M. et al. (2018) Exon Junction Complex Shapes the Transcriptome by Repressing Recursive Splicing. Mol Cell, 72, 496–509 e499.

23. Bao, X., Guo, X., Yin, M., Tariq, M., Lai, Y., Kanwal, S., Zhou, J., Li, N., Lv, Y., Pulido-Quetglas, C. et al. (2018) Capturing the interactome of newly transcribed RNA. Nat Methods, 15, 213–220.

24. Huang, R., Han, M., Meng, L. and Chen, X. (2018) Transcriptome-wide discovery of coding and noncoding RNA-binding proteins. Proc Natl Acad Sci U S A, 115, E3879–E3887.

25. Castello, A., Fischer, B., Frese, C.K., Horos, R., Alleaume, A.M., Foehr, S., Curk, T., Krijgsveld, J. and Hentze, M.W. (2016) Comprehensive Identification of RNA-Binding Domains in Human Cells. Mol Cell, 63, 696–710.

26. Hayashi, Y., Caboni, L., Das, D., Yumoto, F., Clayton, T., Deller, M.C., Nguyen, P., Farr, C.L., Chiu, H.J., Miller, M.D. et al. (2015) Structure-based discovery of NANOG variant with enhanced properties to promote self-renewal and reprogramming of pluripotent stem cells. Proc Natl Acad Sci U S A, 112, 4666–4671.

27. Brodsky, S., Jana, T., Mittelman, K., Chapal, M., Kumar, D.K., Carmi, M. and Barkai, N. (2020) Intrinsically Disordered Regions Direct Transcription Factor In Vivo Binding Specificity. Mol Cell, 79, 459–471 e454.

28. Romero Romero, M.L., Yang, F., Lin, Y.R., Toth-Petroczy, A., Berezovsky, I.N., Goncearenco, A., Yang, W., Wellner, A., Kumar-Deshmukh, F., Sharon, M. et al. (2018) Simple yet functional phosphate-loop proteins. Proc Natl Acad Sci U S A, 115, E11943–E11950.

29. Uversky, V.N. (2013) The alphabet of intrinsic disorder: II. Various roles of glutamic acid in ordered and intrinsically disordered proteins. Intrinsically Disord Proteins, 1, e24684.

30. Kalkan, T. and Smith, A. (2014) Mapping the route from naive pluripotency to lineage specification. Philos Trans R Soc Lond B Biol Sci, 369.

31. Tu, S., Narendra, V., Yamaji, M., Vidal, S.E., Rojas, L.A., Wang, X., Kim, S.Y., Garcia, B.A., Tuschl, T., Stadtfeld, M. et al. (2016) Co-repressor CBFA2T2 regulates pluripotency and germline development. Nature, 534, 387–390.

32. Zhang, J., Ratanasirintrawoot, S., Chandrasekaran, S., Wu, Z., Ficarro, S.B., Yu, C., Ross, C.A., Cacchiarelli, D., Xia, Q., Seligson, M. et al. (2016) LIN28 Regulates Stem Cell Metabolism and Conversion to Primed Pluripotency. Cell Stem Cell, 19, 66–80.

33. Mayer, D., Stadler, M.B., Rittirsch, M., Hess, D., Lukonin, I., Winzi, M., Smith, A., Buchholz, F. and Betschinger, J. (2020) Zfp281 orchestrates interconversion of pluripotent states by engaging Ehmt1 and Zic2. EMBO J, 39, e102591.

34. Faehnle, C.R., Walleshauser, J. and Joshua-Tor, L. (2017) Multi-domain utilization by TUT4 and TUT7 in control of let-7 biogenesis. Nat Struct Mol Biol, 24, 658–665.

35. Seki, Y. (2018) PRDM14 Is a Unique Epigenetic Regulator Stabilizing Transcriptional Networks for Pluripotency. Front Cell Dev Biol, 6, 12.

36. Zeng, Y., Yao, B., Shin, J., Lin, L., Kim, N., Song, Q., Liu, S., Su, Y., Guo, J.U., Huang, L. et al. (2016) Lin28A Binds Active Promoters and Recruits Tet1 to Regulate Gene Expression. Mol Cell, 61, 153–160.

37. Zagore, L.L., Sweet, T.J., Hannigan, M.M., Weyn-Vanhentenryck, S.M., Jobava, R., Hatzoglou, M., Zhang, C. and Licatalosi, D.D. (2018) DAZL Regulates Germ Cell Survival through a Network of PolyA-Proximal mRNA Interactions. Cell Rep, 25, 1225–1240 e1226.

38. Jenkins, H.T., Malkova, B. and Edwards, T.A. (2011) Kinked beta-strands mediate high-affinity recognition of mRNA targets by the germ-cell regulator DAZL. Proc Natl Acad Sci U S A, 108, 18266–18271.

39. Konig, J., Zarnack, K., Rot, G., Curk, T., Kayikci, M., Zupan, B., Turner, D.J., Luscombe, N.M. and Ule, J. (2010) iCLIP reveals the function of hnRNP particles in splicing at individual nucleotide resolution. Nat Struct Mol Biol, 17, 909–915.

40. Rafiee, M.R., Sigismondo, G., Kalxdorf, M., Forster, L., Brugger, B., Bethune, J. and Krijgsveld, J. (2020) Protease-resistant streptavidin for interaction proteomics. Mol Syst Biol, 16, e9370.

41. Hu, G., Wu, Z., Uversky, V.N. and Kurgan, L. (2017) Functional Analysis of Human Hub Proteins and Their Interactors Involved in the Intrinsic Disorder-Enriched Interactions. Int J Mol Sci, 18.

42. Zhu, L. and Brangwynne, C.P. (2015) Nuclear bodies: the emerging biophysics of nucleoplasmic phases. Curr Opin Cell Biol, 34, 23–30.

43. Shin, Y., Chang, Y.C., Lee, D.S.W., Berry, J., Sanders, D.W., Ronceray, P., Wingreen, N.S., Haataja, M. and Brangwynne, C.P. (2019) Liquid Nuclear Condensates Mechanically Sense and Restructure the Genome. Cell, 176, 1518.

44. Wei, M.T., Chang, Y.C., Shimobayashi, S.F., Shin, Y., Strom, A.R. and Brangwynne, C.P. (2020) Nucleated transcriptional condensates amplify gene expression. Nat Cell Biol, 22, 1187–1196.

45. Boija, A., Klein, I.A., Sabari, B.R., Dall’Agnese, A., Coffey, E.L., Zamudio, A.V., Li, C.H., Shrinivas, K., Manteiga, J.C., Hannett, N.M. et al. (2018) Transcription Factors Activate Genes through the Phase-Separation Capacity of Their Activation Domains. Cell, 175, 1842–1855 e1816.

46. Garcia, D.A., Johnson, T.A., Presman, D.M., Fettweis, G., Wagh, K., Rinaldi, L., Stavreva, D.A., Paakinaho, V., Jensen, R.A.M., Mandrup, S. et al. (2021) An intrinsically disordered region-mediated confinement state contributes to the dynamics and function of transcription factors. Mol Cell, 81, 1484–1498 e1486.

47. Li, H., Liang, Z., Yang, J., Wang, D., Wang, H., Zhu, M., Geng, B. and Xu, E.Y. (2019) DAZL is a master translational regulator of murine spermatogenesis. Natl Sci Rev, 6, 455–468.

48. Gibbs, E.B. and Kriwacki, R.W. (2018) Linker histones as liquid-like glue for chromatin. Proc Natl Acad Sci U S A, 115, 11868–11870.

49. Turner, A.L., Watson, M., Wilkins, O.G., Cato, L., Travers, A., Thomas, J.O. and Stott, K. (2018) Highly disordered histone H1-DNA model complexes and their condensates. Proc Natl Acad Sci U S A, 115, 11964–11969.

50. Long, Y., Hwang, T., Gooding, A.R., Goodrich, K.J., Rinn, J.L. and Cech, T.R. (2020) RNA is essential for PRC2 chromatin occupancy and function in human pluripotent stem cells. Nat Genet, 52, 931–938.

51. Ginno, P.A., Burger, L., Seebacher, J., Iesmantavicius, V. and Schubeler, D. (2018) Cell cycle-resolved chromatin proteomics reveals the extent of mitotic preservation of the genomic regulatory landscape. Nat Commun, 9, 4048.

52. Alabert, C., Bukowski-Wills, J.C., Lee, S.B., Kustatscher, G., Nakamura, K., de Lima Alves, F., Menard, P., Mejlvang, J., Rappsilber, J. and Groth, A. (2014) Nascent chromatin capture proteomics determines chromatin dynamics during DNA replication and identifies unknown fork components. Nat Cell Biol, 16, 281–293.

53. Kliszczak, A.E., Rainey, M.D., Harhen, B., Boisvert, F.M. and Santocanale, C. (2011) DNA mediated chromatin pull-down for the study of chromatin replication. Sci Rep, 1, 95.

54. Kohlmeier, F., Maya-Mendoza, A. and Jackson, D.A. (2013) EdU induces DNA damage response and cell death in mESC in culture. Chromosome Res, 21, 87–100.

55. Hoffman, E.A., Frey, B.L., Smith, L.M. and Auble, D.T. (2015) Formaldehyde crosslinking: a tool for the study of chromatin complexes. J Biol Chem, 290, 26404–26411.

