## Supplementary Material with figures for "Chromatin-contact atlas reveals disorder-mediated protein interactions and moonlighting chromatin-associated RBPs"

#### Step by step SPACE procedure

**Required material:**

· Formaldehyde (Methanol-free, Pierce 28906, or 28908)

· BCA protein assay kit (Thermo, 23225)

· Benzonase (Sigma, E8263)

· DNA Binding Beads (Thermo 4489112)

· Guanidinium thiocyanate ( Sigma, G9277-100G)

· N-Lauroylsarcosine sodium salt (Sigma, L9150-100G)

· Tris HCl (Sigma, T2319-1L)

· EDTA (Sigma, 324504-500ML)

· 2-Propanol (Sigma, 278475-1L)

· Ammonium Bicarbonate ( Sigma, 09830-500G)

· DTT (D9779-5G)

· Iodoacetamide (Sigma I1149-5G)

· RNase A (Thermo EN0531)

· Acetonitrile (271004-1L)

· Trypsin (Promega, V5280)

· LysC (Wako, 121-05063)

· Trifluroacetic acid (Sigma 302031)

· Formic acid (Sigma 5.43804)

· ZipTip with 0.6 µL $C_{18}$ resin, (Merck, ZTC18S096)

**Reagents:**

· PBS-T(PBS + Tween 0.1% )

· PBS-SDS (PBS + SDS 1%)

· Elution buffer: Tris HCl 10mM pH 7.5

· Lysis buffer: Guanidinium thiocyanate 4M, Tris HCl 100mM, Sarkosyl 2%, EDTA 10mM

· Wash buffer 1: Lysis buffer + 2-propanol (1:1 v/v)

· Wash buffer 2: Ethanol 80% (v/v)

· AMBIC buffer: Ammonium Bicarbonate 50mM, DTT 10mM (prepare freshly)

· DTT solution: DTT 1M (keep the aliquots in -20 °C, thaw once)

· IAA solution: IAA 0.4M (keep the aliquots in -20 °C, thaw once)

**Equipments:**

· Diagenode Bioruptor Pico

· James Products Ultrasonic 7000S (ultrasonic cleaner)

· Magnetic stand for 15ml tubes

· Magnetic stand for 2ml tubes

· Magnetic stand for PCR tubes

**Experimental procedure:**

A) Formaldehyde crosslinking:

The optimal number of cells for a SPACE experiment is 100,000 – 500,000 cells per replicate. You may use even less than 100,000 cells. However, we don't recommend to use more than 3,000,000 cells.

1. formaldehyde crosslinking can be carried out in the medium of the cells, by adding 16% formaldehyde in the medium of the cells to make 1% final concentration. You may add formaldehyde directly to the medium of the cells in their plates. Obviously, as the non-crosslinked control you don't need to add formaldehyde to the medium.

Note: formaldehyde is toxic, and you should work in a fume hood.

2. Keep the plates 10min in the fume hood.

3. Discard the medium of the cells, as appropriate. Wash the cells two times with PBS.

4. Pour PBS-T in the plates, half of the volume of the medium.

5. Lift the cells by a cell lifter.

6. Collect the cell suspension, and transfer them to a 15-ml/50-ml tube.

7. Repeat steps 4-6 once again to collect most of the cells.

8. Spin the tubes in 400g (2000 RPM in an Eppendorf/Thermo centrifuge) for 2min

9. Discard the supernatant, and freeze the cells in -80 °C

Note: the cells are stable in -80 °C for months.

B) Chromatin purification.

1. Thaw the cells, and resuspend them in PBS-T. The volume depends on the number of the cells. For example you may resuspend the cells in 10% of the original volume of the medium. (i.e. 10 cm dish: 1ml, 6w plate: 0.2ml). The idea is to measure protein concentrations, to start the experiment from equal amounts of input cells.

   Note: Don't use more 3 million cells as the starting material.

2. Take 5-10ul of the samples, and increase the volumes to 30ul by PBS-SDS. Heat the samples at 95 °C for 5 min. After heating the samples, they will become viscous due to the release of DNA. Add 0.5ul Benzonase, and wait a few minutes before proceeding to the next steps. Vortex the samples gently, and measure the protein concentrations by BCA assay. The BSA standards should be diluted in PBS-SDS, and should be heated like the unknown samples.

3. Pour equal amounts of samples in 15-ml tubes. Add 10ul RNase A. Vortex gently, and wait 5 min.

4. Add 2ml Lysis buffer, and vortex the tubes vigorously.

5. Add 2ml 2-propanol, and vortex the tubes vigorously.

6. You may pool SILAC labelled samples. For example, you may pool heavy SILAC crosslinked samples with light SILAC non-crosslinked samples at this step.

Note: if you see white precipitations spin the tube briefly to separate them. Normally it happens in non-crosslinked controls. Do not carry the precipitations to the pool.

7. Add 30ul to 50ul DNA-binding beads to the mixture. It's not necessary to wash or dilute the beads.

8. Vortex the tubes, and wait 10min.

9. Put the tubes on the magnet, and wait 10min to separate the beads.

10. Gently remove the supernatant.

Note: the supernatant is probably brownish even after 10min. It's OK if you lose some beads.

11. Resuspend the beads in 1ml of Wash buffer 1, and transfer them to 2-ml tubes.

12. Vortex the tubes, spin briefly, and put them on the magnet.

13. Wait 2min to separate the beads on the magnet. Discard the supernatant.

14. Remove the beads from the magnet, add 1ml Wash buffer 2.

15. Vortex the tubes, spin briefly, and put them on the magnet.

16. Wait 2min to separate the beads on the magnet. Discard the supernatant.

17. Vortex the tubes, spin briefly, and put them on the magnet.

18. Discard the residual Wash buffer 2.

19. Add 200ul of elution buffer. Do not pipette as the beads are very sticky at this point.

20. Put the tube in the ultrasonic cleaner, and turn on the sonicate them for 5min.

21. Transfer the tubes to the rotor of Diagenode Bioruptor.

22. Sonicate the tubes 3 cycles: 30 sec ON, 30 sec OFF

    Note: the beads should be resuspended at this step.

23. Add 10ul of RNase A (10 mg/ml) to the samples, and agitate them (1000 RPM) in a thermomixer at 37 °C for 15 min

24. Add 250ul of Lysis buffer, and vortex the tubes.

25. Add 300ul of 2-Propanol, and vortex the tubes.

26. Wait 10min.

27. Put the tubes on the magnet, and wait 10min to separate the beads.

28. Gently remove the supernatant.

29. Remove the beads from the magnet, add 1ml Wash buffer 2.

30. Vortex the tubes, spin briefly, and put them on the magnet. Discard the wash buffer.

31. Repeat the last 2 steps once again.

32. Remove the beads from the magnet, add 0.5ml Acetonitrile 100%

33. Vortex the tubes, spin briefly, and put them on the magnet.

34. Resuspend the beads in 80ul of Acetonitrile 100%, and transfer them to PCR tubes.

Note: If you use > 1 million cells you may need to cut the head of the 200-ul tips to be able to transfer the beads.

You may repeat this step once again to transfer all the beads to a PCR tube.

35. Put the cells on a magnet, and discard the supernatant.

36. Add 18ul of AMBIC buffer (if you wish to use TMT labelling use TEAB buffer)

37. Heat the tubes in a PCR machine at 95 °C for 10min. Turn on the lid (105 °C ) to avoid evaporation.

38. Chill the tubes on ice. Add 2ul of the IAA solution. Vortex gently, and keep in a drawer for 15min.

39. Add 1ul of the DTT solution, and vortex gently.

40. Add 400ng trypsin, 100ng LysC, and incubate the tubes 12-14 h at 37 °C. Turn on the lid (80 °C) to avoid evaporation.

41. Add 100ng trypsin, vortex and continue the digestion for another 2-h.

42. Remove the beads on the magnet, and transfer the supernatant to a new PCR tube

43. Clean-up the peptides using stage-tips or ZipTips.

### Step by step SPACE-SICAP procedure

**Additional required material:**

· Biotin-ddUTP (Jenabioscience, NU-1619-BIOX-S or L)

· Biotin-dCTP (Jenabioscience, NU-809-BIOX-S)

· Streptavidin magnetic beads (NEB S1420S)

· Sodium cyanoborohydride (Sigma, 8180530025)

· TdT (Thermo Scientific, EP0162)

· T4 PNK (NEB, M0201S)

· Amicon Ultra-0.5 Centrifugal Filter Unit, 30 KDa (Millipore, UFC503096)

· SPRIselect (Beckman Coulter, B23317)

**Reagent preparation:**

· Protease-resistant streptavidin beads: Beads should be treated in fume hood. For 5ml beads, prepare 5ml Sodium cyanoborohydride 0.2M (Reagent A) and 5ml Formaldehyde 4% (Reagent B). Pour the beads in a 15-ml tube, put the tube on the magnet, remove the beads, and discard the supernatant. Wash the beads with 5ml PBS-T, put the tube on the magnet, remove the beads, and discard the supernatant. Add reagent A and reagent B to the beads. Resuspend the beads, and keep them in the hood for 2 hours with occasional mixing. Put the tube on the magnet, remove the beads, and discard the supernatant appropriately. Wash the beads with 5ml Tris HCl 0.1 M pH 7.5, twice. Finally, resuspend the beads in 5ml PBS-T, and keep them in the fridge. The beads are now resistant to Lys-C digestion, and they are stable for months. For more details, please refer to (Rafiee et al., MSB, 2020)

**Experimental procedure:**

The required number of cells for a SPACE-SICAP experiment is 2.5 million cells per replicate.

A) DNA labelling and chromatin purification

1. After RNase A treatment in SPACE procedure (Step 22, section B), remove the beads on the magnet. Transfer the supernatant to Amicon ultrafilteration tubes, and spin them in 12000g for 7 min at 8 °C. The volume of the liquid in the column should be < 100ul, otherwise, continue the centrifuge for a few more minutes.

2. Collect the liquid in the column, and transfer them to PCR tubes.

3. Add the following reagents:

    I. TdT buffer 20ul

    II. Biotin-ddUTP 5ul

    III. Biotin-dCTP 5ul

    IV. TdT 3ul

    V. T4 PNK 1ul

    VI. H2O fill up to 100ul

4. Vortex, and spin briefly

5. Incubate the tubes at 37 °C for 30min

6. Add 100ul SPRIselect beads, vortex, and spin briefly

7. Wait 10min

8. Separate the beads on the magnet

9. Wash the beads with ethanol 80% without disturbing the beads

10. Repeat the last step once again

11. Remove the residual ethanol

12. Resuspend the beads in 100ul elution buffer

13. Sonicate the tubes in the ultrasonic cleaner for 5min

14. Remove the beads on the magnet, and transfer the supernatant to 2ml tubes

15. Increase the volume to 1ml with PBS-SDS

16. Add 75ul protease-resistant streptavidin beads.

17. Rotate 45min at room temperature

18. Separate the beads on the magnet, and discard the supernatant

19. Wash the beads two times with PBS-SDS.

Note: during this step you will observe the beads are dispersed on the magnetic stand

20. Wash the beads with 2-propanol 20%

21. Wash the beads two times with Acetonitrile 40%

22. Separate the beads on the magnet, and discard the supernatant

23. Resuspend the beads in 80ul of Acetonitrile 40%, and transfer them to PCR tubes.

Note: You may also need to repeat this step once again to transfer all the beads to a PCR tube.

24. Put the cells on a magnet, and discard the supernatant.

25. Add 18ul of AMBIC buffer (if you wish to use TMT labelling use TEAB buffer)

26. Heat the tubes in a PCR machine at 50 °C for 15min. Turn on the lid (80 °C ) to avoid evaporation.

27. Chill the tubes on ice. Add 2ul of the IAA solution. Vortex gently, and keep in a drawer for 15min.

28. Add 1ul of the DTT solution, and vortex gently.

29. Add 400ng LysC, and incubate the tubes 12-14 h at 37 °C. Turn on the lid (80 °C) to avoid evaporation.

30. Separate the beads on the magnet. Transfer the supernatant to new PCR tubes.

31. Add 300ng trypsin and continue the digestion for 4-8 hours.

32. Clean-up the peptides using stage-tips or ZipTips.

#### Step by step SPACEmap

After step 36 of SPACE (Pour 18ul AMBIC on the beads)

1. Heat the tubes in a PCR machine at 50 °C for 10min. Turn on the lid (80 °C ) to avoid evaporation.
2. Chill the tubes on ice. Add 2ul of the IAA solution. Vortex gently, and keep in a drawer for 15min.
3. Add 0.5ul of 1M DTT solution, and vortex gently.
4. Add 1000ng Trypsin, 200ng LysC, and incubate the tubes 12-16 h at 37 °C. Turn on the lid (80 °C) to avoid evaporation.
5. Add 500ng Trypsin, and incubate the tubes 4 h at 37 °C.
6. Add 80ul SPACE Lysis buffer, and vortex
7. Add 80ul 2-propanol 100%, and vortex
8. Wait 10min, and spin the tubes briefly
9. Separate the beads on magnet
10. Collect the supernatant, and transfer it to a 1.5ml tube. This is the **Released fraction** to be compared with the crosslinked fraction.
11. Wash the DNA-binding beads with Wash buffer 1 once, and discard it
12. Wash the DNA-binding beads with Wash buffer 2 once, and discard it
13. Wash the DNA-binding beads with acetonitrile twice, and discard it
14. Briefly spin the beads, and discard the residues of acetonitrile in the bottom of the tube
15. Resuspend the beads in 20ul of AMBIC buffer
16. Heat the tubes in a PCR machine at 95 °C for 10min. Turn on the lid (105 °C ) to avoid evaporation.
17. Chill the samples, and add 300ng trypsin, 50ng lysC, and incubate the tubes 12 h at 37 °C. Turn on the lid (80 °C) to avoid evaporation.
18. Remove the beads on the magnet, and transfer the supernatant to a new PCR tube. This is the **Crosslinked fraction**, and should be compared with the Released fraction to determine the peptides enriched/depleted.
19. Clean-up the peptides using stage-tips or ZipTips.

**Step by step ChIP-SPACE**

After the ChIP protocol, and regular washing steps perform the following steps:

1. Separate the beads on the magnet, and discard the last wash buffer
2. Resuspend the beads in 100ul TE buffer (or Elution buffer), add 5ul RNase A (10mg/ml), and agitate at 750 RPM in a ThermoMixer at 37 °C for 10min.

   Note: Skip this step if you wish to have a RNase negative control.
3. Resuspend the beads in 300ul Lysis buffer, vortex vigorously, and incubate at 37 °C for 2min in a Thermomixer with 1000 RPM agitation.
4. Remove the beads on the magnet, and transfer the supernatant to a new 2-ml tube
5. Add 300ul 2-propanol, and vortex
6. Add 30ul of DNA-binding beads, vortex, and spin
7. Wait 10min
8. Separate the beads on the magnet, and discard the supernatant
9. Wash the beads with 500ul Wash buffer 1, separate the beads on the magnet, and discard the supernatant.
10. Wash the beads with 500ul Wash buffer 2, separate the beads on the magnet, and discard the supernatant.
11. Repeat the last step once again.
12. Wash the beads with 500ul Acetonitrile 100%, separate the beads on the magnet, and discard the supernatant.
13. Resuspend the beads in 80ul of Acetonitrile 100%, and transfer them to PCR tubes.

    Note: You may also need to repeat this step once again to transfer all the beads to a PCR tube.
14. Put the cells on a magnet, and discard the supernatant.
15. Add 18ul of AMBIC buffer (if you wish to use TMT labelling use TEAB buffer)
16. Heat the tubes in a PCR machine at 95 °C for 10min. Turn on the lid (105 °C ) to avoid evaporation.
17. Chill the tubes on ice. Add 2ul of the IAA solution. Vortex gently, and keep in a drawer for 15min.

18. Add 1ul of the DTT solution, and vortex gently.
19. Add 250ng trypsin, 100ng LysC, and incubate the tubes 14-16 h at 37 °C. Turn on the lid (80 °C) to avoid evaporation.
20. Remove the beads on the magnet, and transfer the supernatant to a new PCR tube
21. Clean-up the peptides using stage-tips or ZipTips.

**ChIP-seq and data analysis in detail**

We followed ENCODE guidelines for ChIP-seq. The mESCs were grown in 2iL medium. The cells were detached and fixed by 1.5% formaldehyde in PBS for 15min. About 20 million cells were used for each ChIP experiment. Two replicates were prepared for Dazl ChIP-seq. The cells were resuspended in Tris.HCl pH 8, 10mM, and kept on ice for 5min. Then 1% Triton X100 was added to the cells, and the cell lysate was kept on ice for another 5min. The samples were spun at 2000g for 2min to remove the supernatant. The cells pellets were resuspended in Tris.HCl pH 8, 10mM, and spun at 2000g for 2 min. The cell pellet was resuspended in LB3 buffer (Tris.HCl pH 8 10 mM, NaCl 100 mM, EDTA 1 mM, EGTA 0.5 mM, Na-deoxycholate 0.1%, and N-lauroylsarcosine 0.5%). Chromatin was solubilized by Bioruptor Pico for 10 cycles: 30s on and 30s off. The sheared chromatin was spun at 12000g for 10min to remove the cell debris. The chromatin was sheared to < 500 bp fragments with the peaks about 200-300 bp. Dazl immunoprecipitation was carried out using CST antibody #8042 overnight at 4 °C. The antibody was diluted 1:100 in sheared chromatin. As the negative control, we used 1% of the input. We verified the Dazl antibody using western blotting and mass spectrometry. Following the immunoprecipitation using protein A magnetic beads, the beads were washed 6 times using Tris.HCl 50mM, EDTA 5mM, Triton X100 1% and NP-40 0.5%. Then chromatin fragments were eluted using SDS 2%. Both IP samples and the input control were treated with proteinase K to degrade the proteins. DNA was purified by SPRI beads. The purified DNA was end-repaired by T4 DNA polymerase, Klenow fragment and T4 polynucleotide kinase. Then DNA was purified and subjected to A-tailing by Klenow 3'-exo$^{-}$. DNA was purified and adaptors for illumina sequencing were ligated to the DNA fragments. The libraries were amplified using NEB primer indices and Q5 DNA polymerase for 12 cycles. The library concentrations were determined using the Qubit DNA high sensitivity kit to be 0.662, 0.994 and 56 ng/ul for Dazl_R1, Dazl_R2 and the input control, respectively. The libraries fragment size were checked by TapeStation, and the peaks were determined to be about 300-350 bp. Sequencing was carried out using 100nt reads on paired-end mode by HiSeq4000. Sequencing data quality was evaluated using FastQC package. The total number of reads were about 28million, 20milion and 30 million reads for Dazl_R1, Dazl_R2 and the input control, respectively. The rate of the duplicated reads was determined to be about 48%, 52% and 25% for Dazl_R1, Dazl_R2 and the input control, respectively. The reads were trimmed and aligned to the mouse genome (mm10) using Bowtie2. More than 99% of the reads in all libraries were aligned to the mouse genome. The reads were deduplicated with samtools. ChIP quality was estimated by cross-correlation using the "SPP" tool as suggested by ENCODE ChIP-seq guidelines. Peak calling was performed using MACS2. Reproducibility of the ChIP replicates and final peak selection was carried out using the IDR pipeline at a 1% IDR cutoff for the final list of the peaks. Dazl peaks annotation into genomic features was done using ChIPseeker R package with 3kb around TSS set for promoter region window.

**Domain analysis in detail**

First, we found domains and intrinsically disordered regions (IDRs) in the proteins from the crosslinked and released SPACEmap fractions (Supplementary Figure 4F):

1. We obtained protein sequences from UniProt using UniProt IDs and scanned the sequences for domains and IDRs with InterProScan v5.47-82.0.
2. From the scanning results, we selected matches that have an InterPro ID (for domains) or a MobiDBLite ID (for IDRs).
3. We excluded matches that have an InterPro ID but still do not represent domains or IDRs (ProSitePatterns, PRINTS, PIRSF and PANTHER signatures).
4. We merged highly overlapping matches in the following way. In each InterPro entry, we selected all pairs of matches where matches overlapped by at least 70% of the length of either match. We merged matches in such pairs, obtaining combined matches. We further merged original and combined matches using the same overlap criterion until there were no pairs of matches that met it.
5. We defined the consensus match coordinates as the coordinates of the resulting combined matches and the coordinates of the original matches that did not overlap with any other matches. This allowed us to delineate multiple domain matches within InterPro entries and to merge highly overlapping predicted IDRs. We assigned each consensus match with a corresponding InterPro ID.

Next, we found peptides from the crosslinked and released SPACEmap fractions that matched the domains and IDRs defined in the previous step (Supplementary Figure 4G):

1. We postulated that a domain or an IDR matches a peptide if it overlaps with the peptide or resides no farther than 10 amino acids from the ends of the peptide.
2. Using this definition and the consensus match coordinates obtained in the previous step, we selected domains and IDRs that matched peptides.

Finally, we clustered domains that matched peptides from the crosslinked fraction to obtain more general domain types (Supplementary Figure 4H; we did not do this for the released fraction):

1. We grouped all IDRs into one cluster.
2. We started clustering domains by putting each InterPro ID in its own cluster.
3. If matches of two domains (InterPro IDs) overlapped in at least one protein and in at least 70% of the overlaps the overlap size was at least 70% of both domain lengths, then we merged the clusters where the two domains belonged.

This procedure allowed us to group InterPro entries that matched similar sites and hence described similar domains. We named the clusters with the highest total frequency of matches based on the descriptions of the InterPro IDs that the clusters contained.

### Supplementary Figure legends

**Supplementary Figure 1: SPACE and SPACE-SICAP.** (A) After the SPACE procedure, the proteins were separated using an SDS-PAGE. From the ladder, the first lane shows proteins in the lysis buffer (step 10), the 1$^{st}$ wash (step 13), the 2$^{nd}$ wash (step 16), the 3$^{rd}$ wash (step 28), the 4$^{th}$ wash 4 (step 30), the 5$^{th}$ wash (step 31), non-crosslinked control and the crosslinked sample. (B) SPACE-SICAP procedure. Following step3, chromatin is eluted by sonication, and treated with RNase A. Then silica magnetic beads are removed, and DNA is end-labelled by TdT and biotin-ddUTP or biotin-dCTP. Step 4: chromatin is re-captured on protease-resistant streptavidin beads. Step 5: chromatin-associated proteins are digested to be identified by mass spec.

**Supplementary Figure 2: Characterization of proteins enriched by SPACE and SPACE-SICAP.** (A) The enriched proteins by SPACE were categorized based on their biological processes and molecular functions. The node colours indicate if the proteins are 'known DNA/chromatin binding proteins' (dark green), 'Present in Nucleus' (pale green) or 'unexpected' (yellow) in chromatin composition. The enriched proteins are classified using the following keywords: 1- chromatin remodelling 2- chromatin modification, chromatin modifier, nucleosome assembly, chromatin silencing, histone methylation, chromatin assembly 3- DNA replication, DNA repair, double-strand break repair, DNA damage response, DNA helicase, DNA/RNA helicase, response to DNA damage, telomere maintenance 4- chromosome segregation, chromosome separation, condensed chromosome kinetochore, sister chromatid biorientation 5- transcription factor activity, transcription co-activator, transcription coactivator, transcription corepressor, transcriptional activator, transcriptional repressor or transcription factor binding 6- regulation of DNA-templated transcription, transcription DNA-templated, regulation of transcription, transcription by RNA polymerase II, transcription regulation, RNA polymerase II transcription, transcription from RNA polymerase II, transcription-dependent tethering of RNA polymerase II, rRNA transcription 7- posttranscription, post-transcription 8- cell cycle 9- splicing, spliceosome 10- RNA processing, rRNA processing, polyadenylation, RNA 3'-end, RNA cleavage, RNA modification, RNA capping, RNA modification 11- RNA transport. Among the rest of the proteins, we looked for the following keywords: 1- translation, 2- metabolic process, oxidase, 3- cell adhesion, 4- protein folding, and 5-protein transport. Proteins involved in pluripotency, embryonic stem cell processes and polycomb-group components are shown separately. (B) Frequency of the proteins with basic aminoacid compositional bias and IDRs were compared between total proteome and SPACE proteins. Fisher's exact test was used to compare the differences statistically. * p-value $\leq 0.05$, ** p-value $\leq 0.01$ and *** p-value $< 0.001$. (C) The volcano-plot shows proteins enriched via SPACE-SICAP. Very significantly enriched proteins (adj. p-value $< 0.01$ and log2FC >1), significantly enriched proteins (adj. p-value $< 0.1$ and log2FC >1), and proteins that not significantly enriched were shown by dark blue, blue, and grey, respectively. (D) The Venn diagram indicates the number of identified proteins by SPACE-SICAP in comparison to the DNase-treated sample. (E) The Venn diagram shows the intersection of SPACE and SPACE-SICAP. The proteins were categorized into the three groups: known DNA/chromatin binding proteins' (dark green), 'Present in Nucleus' (pale green) or 'unexpected' (yellow). (F) Titration of the input cell number for SPACE procedure.

**Supplementary Figure 3: SPACE and SPACE-SICAP reveal RBPs as a major component of chromatome.** (A) Comparing proteins identified by SPACE (enriched or unenriched) with RICK and CARIC datasets. (B) the Venn diagram shows the overlap of SPACE and SPACE-SICAP proteins. The bar shows frequency of RBPs in the intersect of the datasets. (C) The bars show frequency of RBPs in SPACE-SICAP based on protein counts (left) and relative iBAQ (right).

**Supplementary Figure 4: Chromatin binding sites of the proteins revealed by SPACEmap.** (A) Product of SPACEmap after the first tryptic digestion. Most of the proteins are digested to short peptides by trypsin. Only peptides directly crosslinked to DNA (red) or indirectly via another crosslinked peptide (orange) remain after the digestion. (B-C) Prediction of intrinsically disordered regions in Nanog and Sox2 using IUPred2 (57). The red boxes show peptides enriched in the crosslinked fraction. The pink boxes show DNA-binding domains. (D) Top 10 domains/regions by the proportion of RBPs that contain them (left) or by the proportion of crosslinked fraction peptides of RBP origin that contain them (right). Crosslinked fraction peptides containing these domains/regions overlap with them or reside no farther than 10 amino acids from them. (E) Top 10 domains/regions by the proportion of DNA/chromatin-binders that contain them (left) or by the proportion of crosslinked fraction peptides of DNA/chromatin-binders that contain them (right). Crosslinked fraction peptides containing these domains/regions overlap with them or reside no farther than 10 amino acids from them. (F) Finding matches of domains and intrinsically disordered regions (IDRs) in proteins from the crosslinked and released SPACEmap fractions. We scanned a protein sequence using InterProScan and then excluded non-domain matches while retaining domains and IDRs. Next, we merged highly overlapping matches and defined consensus match coordinates (red triangles denote consensus match starts, blue triangles denote consensus match stops). (G) Selecting domains and IDRs that match peptides from the crosslinked or released SPACEmap fraction. We postulate that a domain or an IDR matches a peptide if it overlaps with the peptide or resides no farther than 10 amino acids from the ends of the peptide. Using this definition and consensus match coordinates of domains and IDRs, we selected only those domains and IDRs that matched peptides. (H) A schematic illustrating clustarisation of similar domains (InterPro entries) that match peptides from the crosslinked SPACEmap fraction. Suppose, we have five proteins whose crosslinked peptides matched some domains and IDRs (left part of the panel). We denote matched IDRs by black rectangles and matched domains by rectangles of different other colours (domain A is denoted by the red rectangle, B – by the blue rectangle, C – by the orange rectangles, D – by the purple rectangles, E – by the green rectangles, and F – by the cyan rectangles). We start by putting all IDRs into one cluster denoted by the framed black label "IDRs" and each of the domains A-F into their own clusters denoted by the framed labels "A"-"F" of the corresponding colours (see the right part of the panel). Next, we check each pair of domains that overlap in at least one protein and put the domains in the same cluster if in at least 70% of the overlaps the size of the overlap is at least 70% of the lengths of both domains. In this schematic, there is only one overlap between domains A and B, and its length comprises at least 70% of the lengths of both domains. Hence, we put domains A and B in the same cluster by merging individual clusters "A" and "B" (see on the right). Next, there are four overlaps between domains D and E, and three of them (i. e., 75% of all D-E overlaps) have lengths that comprise at least 70% of the lengths of both domains. Hence, we put domains D and E in the same cluster by merging individual clusters "D" and "E". Finally, there are two overlaps between domains E and F, and the lengths of both overlaps comprise at least 70% of the lengths of both domains. Hence, we put domains F and E in the same cluster by merging the individual cluster "F" with the cluster that contains the domain E (i. e., the cluster "D, E"). There are no other overlapping pairs of domains, so the final set of domain clusters is {"A, B"; "C"; "D, E, F"}. Hence, instead of 6 different domains (InterPro entries), we obtained 3 general domain types represented by the 3 domain clusters. IDRs comprised a separate cluster from the start of the procedure.

**Supplementary Figure 5: Comparing 2iL and serum states of mES cells using SPACE.**

(A) Comparing proteins identified by SPACE and total proteome analysis of mES cells in 2iL and serum conditions. (B) The differentially regulated proteins in the intersection of the total proteome and SPACE were compared. The boxes indicate the inter-quantile regions (IQR). The line in the box is median. The whiskers show 1.5 IQR. The spots are outliers. (C) Comparative SPACE between mES cells in 2iL and serum condition. Purple points show RBPs with differential binding ratios to chromatin in 2i and serum conditions. (D) SPACE/total proteome iBAQ ratios was calculated, and the range of min to max was divided into 4 classes. This is a replica of Figure 2E to highlight Dazl SPACE/total proteome ratio. The frequency of 1- 'known DNA/chromatin-binders' (dark green), 2- 'proteins present in nucleus' (pale green) and 3- 'unexpected' proteins (yellow), was shown in each class.

**Supplementary Figure 6: Validation of Dazl as a caRBP in mES cells.**

(A) Confocal fluorescence microscopy imaging of fixed mouse embryonic stem cells. Staining was carried out using DAPI, Suz12 and Dazl antibodies. Fluorophore exclusion from subnuclear structures (putative nucleoli) was observed in Dazl staining corroborating its nuclear presence (arrow shows one representative example). (B) ChIP-seq peaks indicate Dazl binds to the promoter of several Hox genes. (C) ChIP-SPACE procedure: after the ChIP procedure (1), samples are treated with and without RNase A (2), then SPACE beads are added to purify chromatin (3), following mass spectrometry (4) proteins co-localized with the target protein are identified. In addition, proteins sensitive to RNase treatment are determined.

### Suppl. Figure 1

A **SPACE on SDS-PAGE**

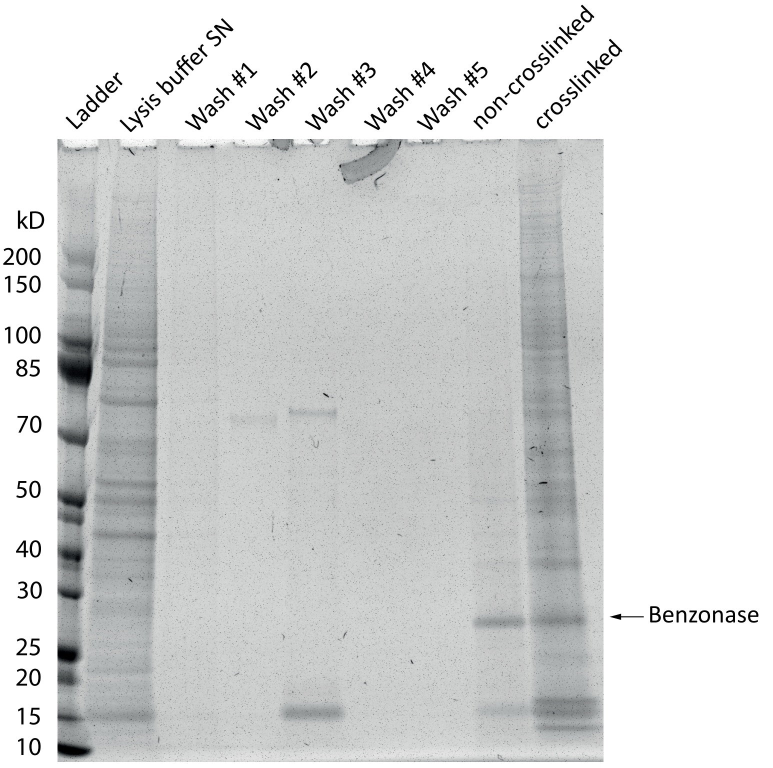


B **SPACE-SICAP**

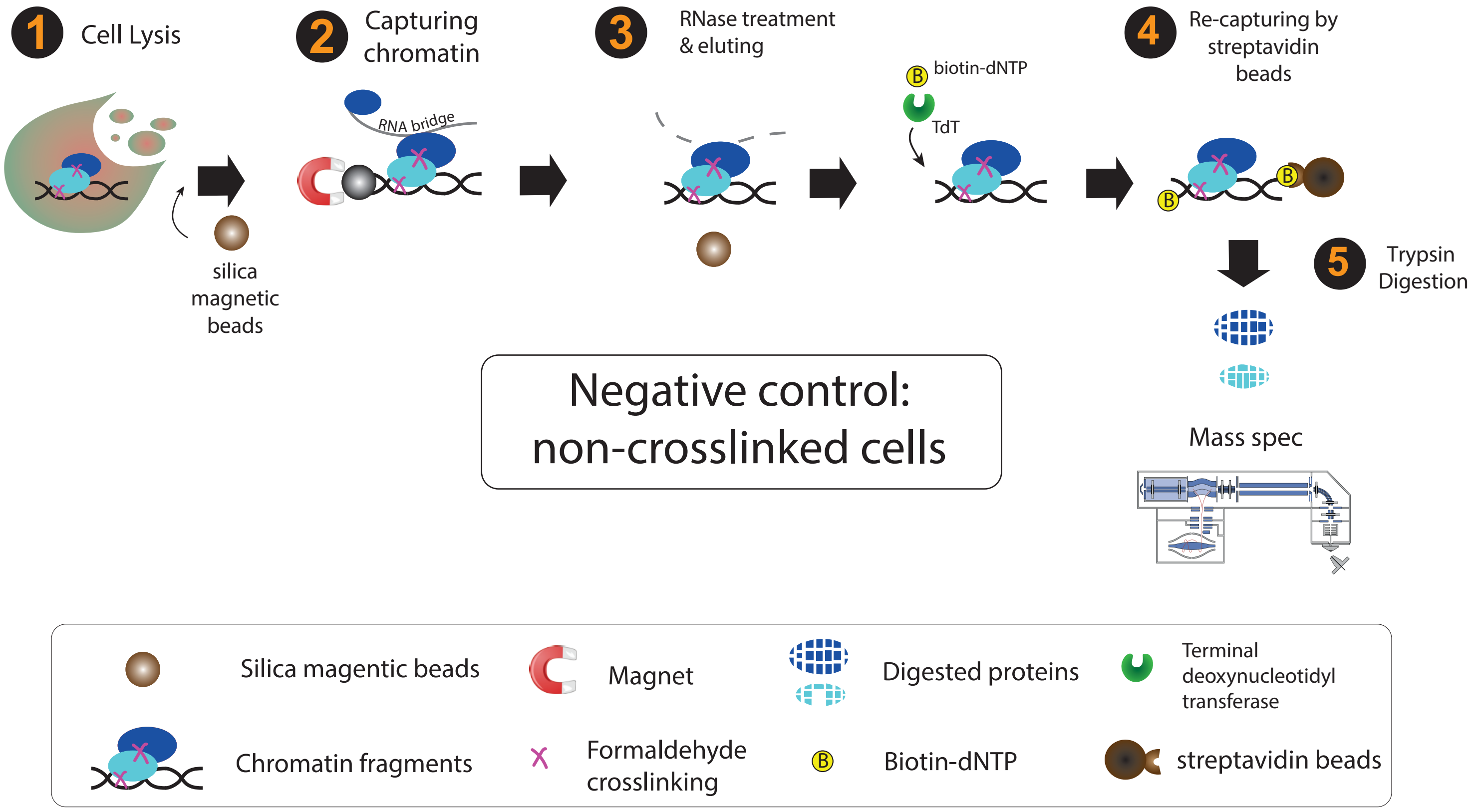

Suppl. Figure 2

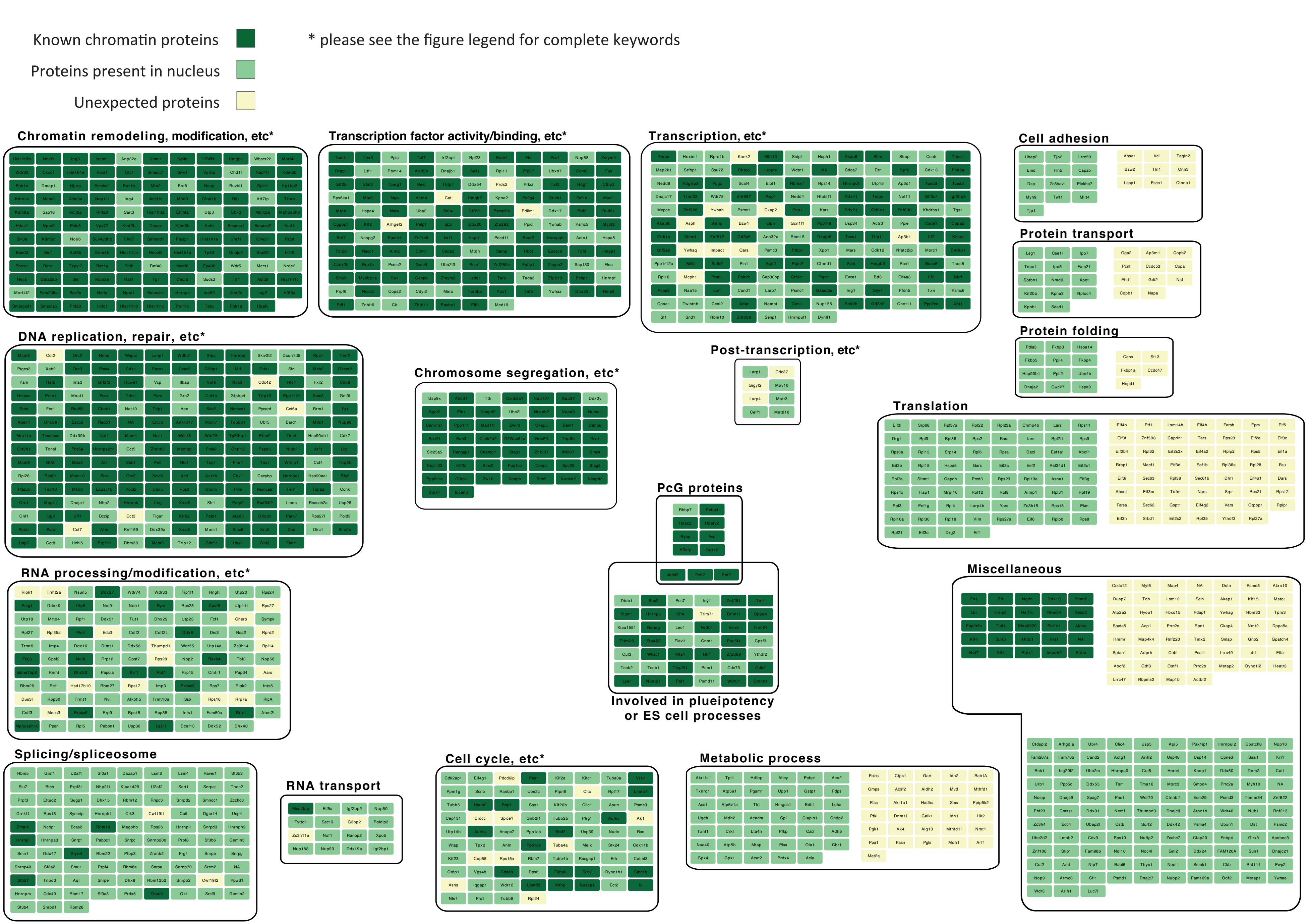

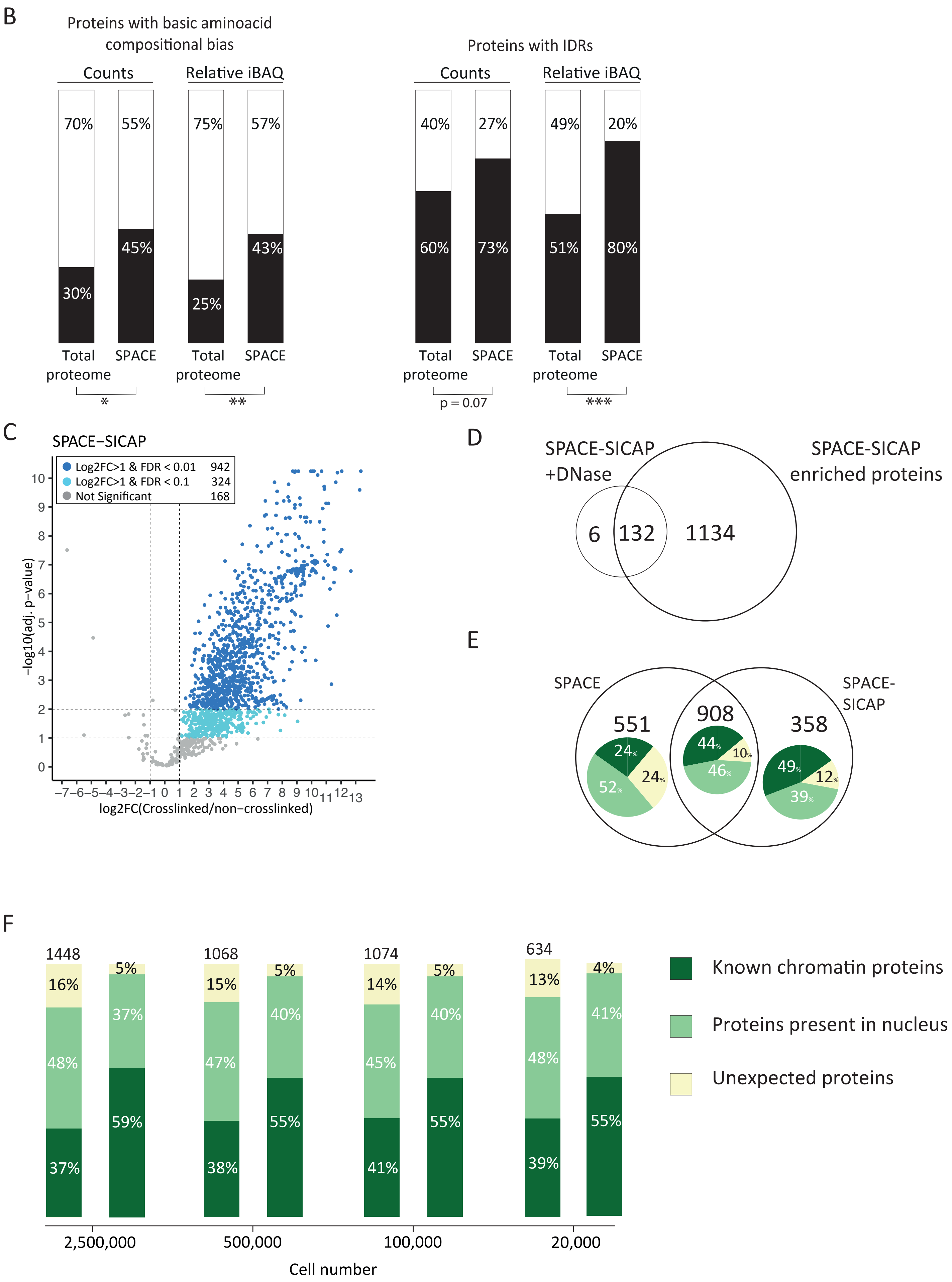
B
Proteins with basic aminoacid compositional bias
Counts
Relative iBAQ
70%
55%
75%
57%
45%
43%
30%
25%
Total proteome
SPACE
Total proteome
SPACE
*
**
Proteins with IDRs
Counts
Relative iBAQ
40%
27%
49%
20%
60%
73%
51%
80%
Total proteome
SPACE
Total proteome
SPACE
p = 0.07
***
C
SPACE−SICAP
Log2FC>1 & FDR < 0.01 942
Log2FC>1 & FDR < 0.1 324
Not Significant 168
−log10(adj. p−value)
log2FC(Crosslinked/non−crosslinked)
D
SPACE-SICAP +DNase
SPACE-SICAP enriched proteins
6
132
1134
E
SPACE
SPACE-SICAP
551
908
358
24%
24%
52%
44%
10%
46%
49%
12%
39%
F
1448
1068
1074
634
16%
48%
37%
5%
37%
59%
15%
47%
38%
5%
40%
55%
14%
45%
41%
5%
40%
55%
13%
48%
39%
4%
41%
55%
2,500,000
500,000
100,000
20,000
Cell number
Known chromatin proteins
Proteins present in nucleus
Unexpected proteins

### Suppl. Figure 3

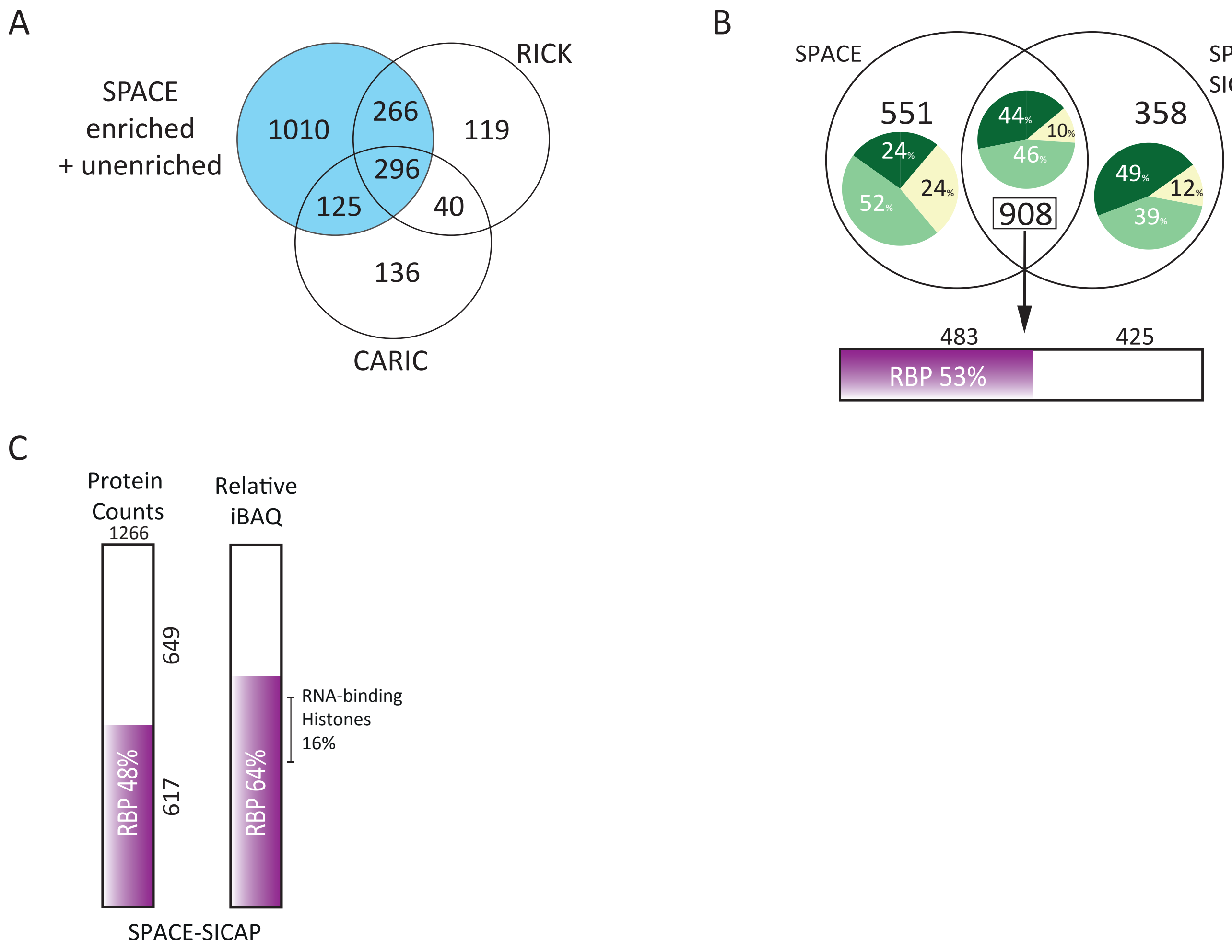

### Suppl. Figure 4

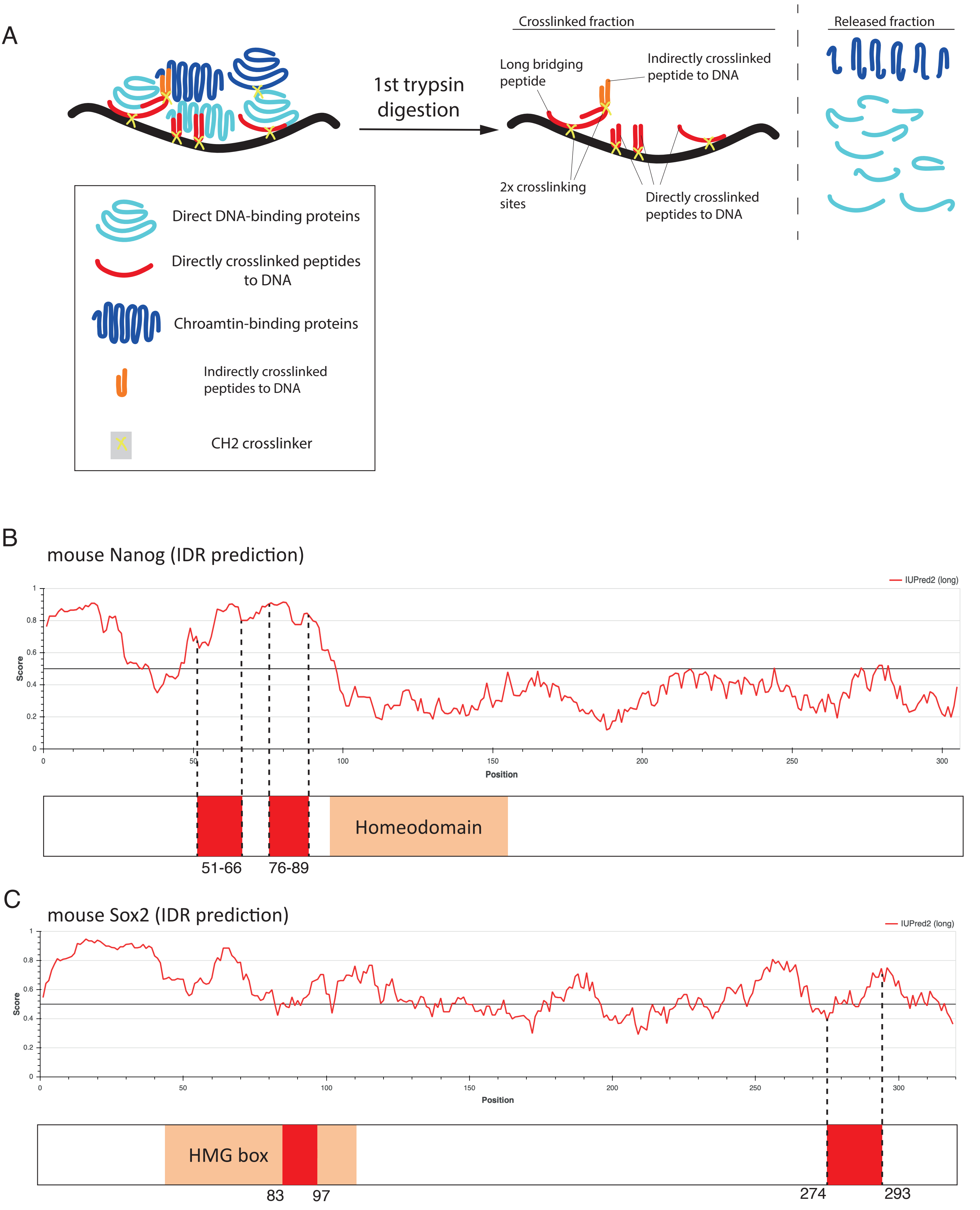

D

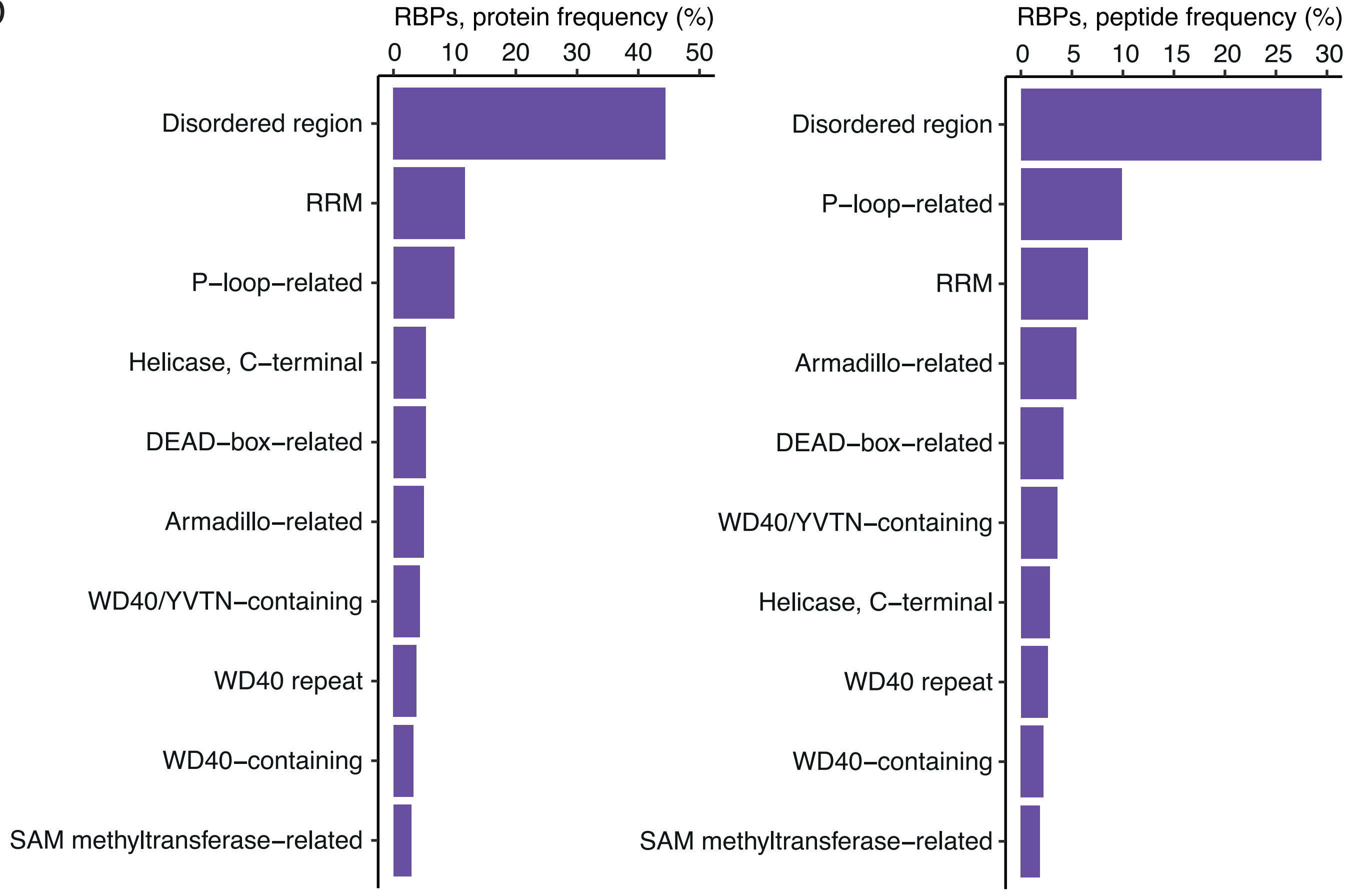
RBPs, protein frequency (%)
0
10
20
30
40
50
Disordered region
RRM
P–loop–related
Helicase, C–terminal
DEAD–box–related
Armadillo–related
WD40/YVTN–containing
WD40 repeat
WD40–containing
SAM methyltransferase–related
RBPs, peptide frequency (%)
0
5
10
15
20
25
30
Disordered region
P–loop–related
RRM
Armadillo–related
DEAD–box–related
WD40/YVTN–containing
Helicase, C–terminal
WD40 repeat
WD40–containing
SAM methyltransferase–related

E

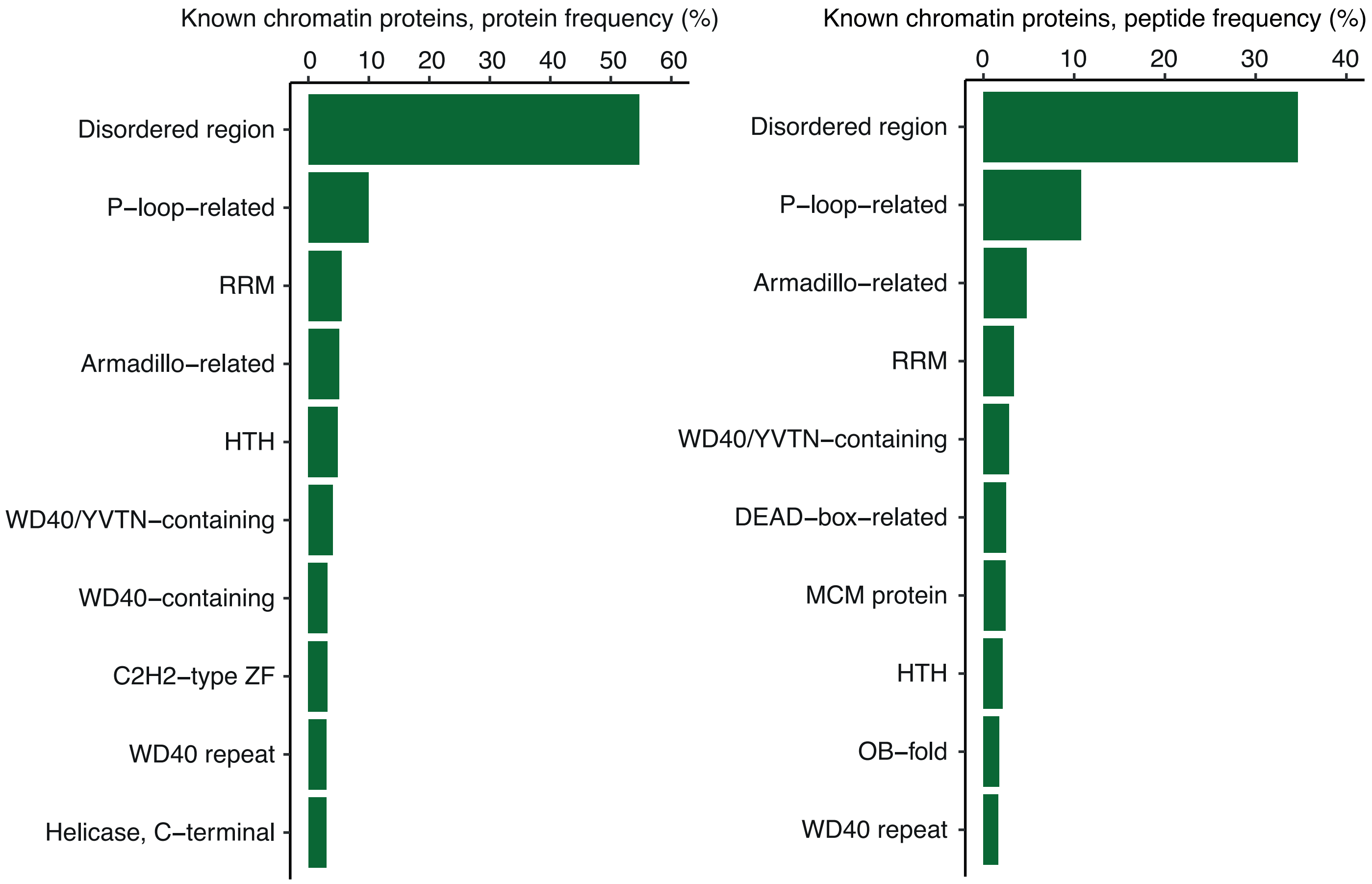
Known chromatin proteins, protein frequency (%)
0
10
20
30
40
50
60
Disordered region
P–loop–related
RRM
Armadillo–related
HTH
WD40/YVTN–containing
WD40–containing
C2H2–type ZF
WD40 repeat
Helicase, C–terminal
Known chromatin proteins, peptide frequency (%)
0
10
20
30
40
Disordered region
P–loop–related
Armadillo–related
RRM
WD40/YVTN–containing
DEAD–box–related
MCM protein
HTH
OB–fold
WD40 repeat

F

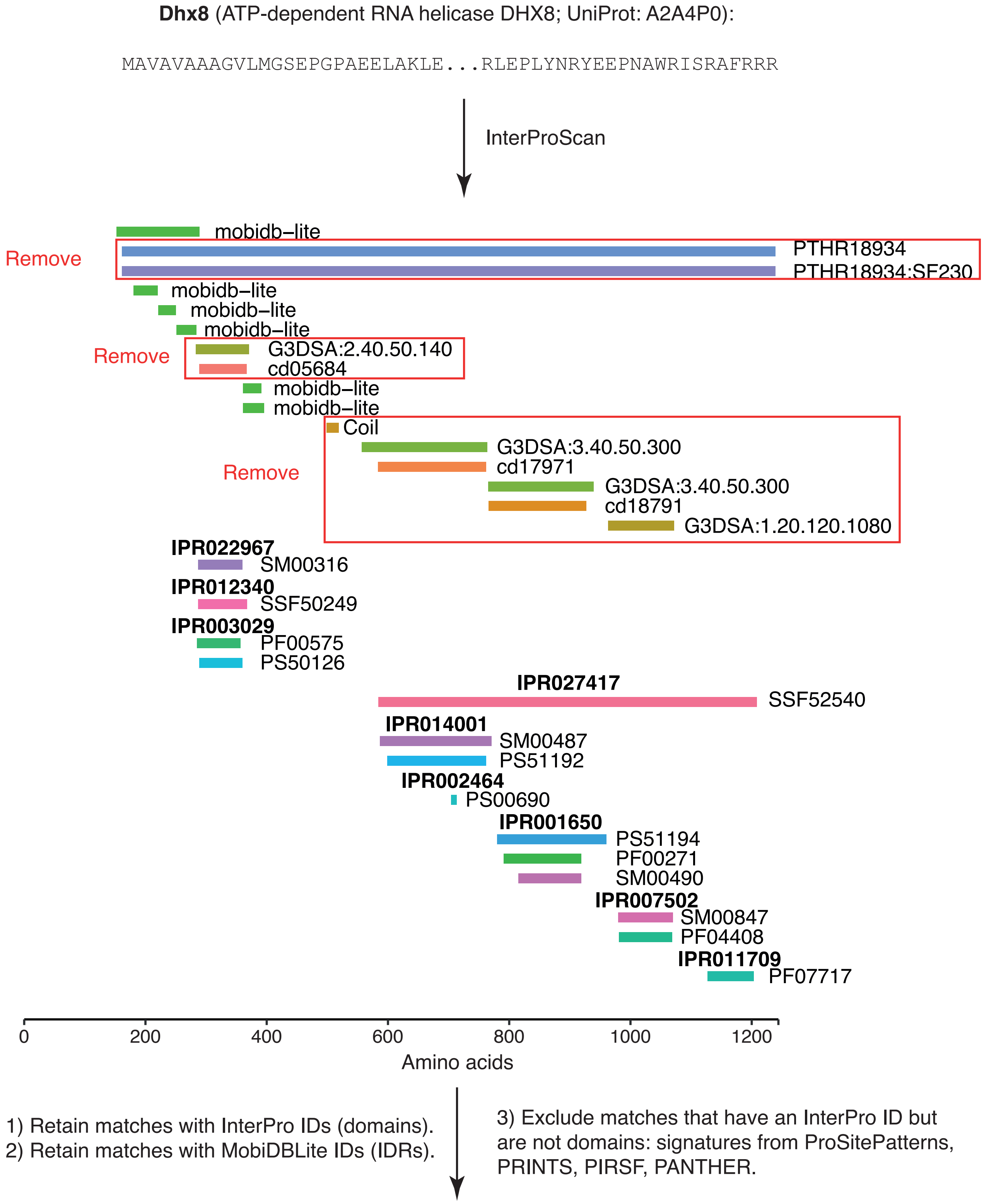


(Continued on the next page.)

F (continued)

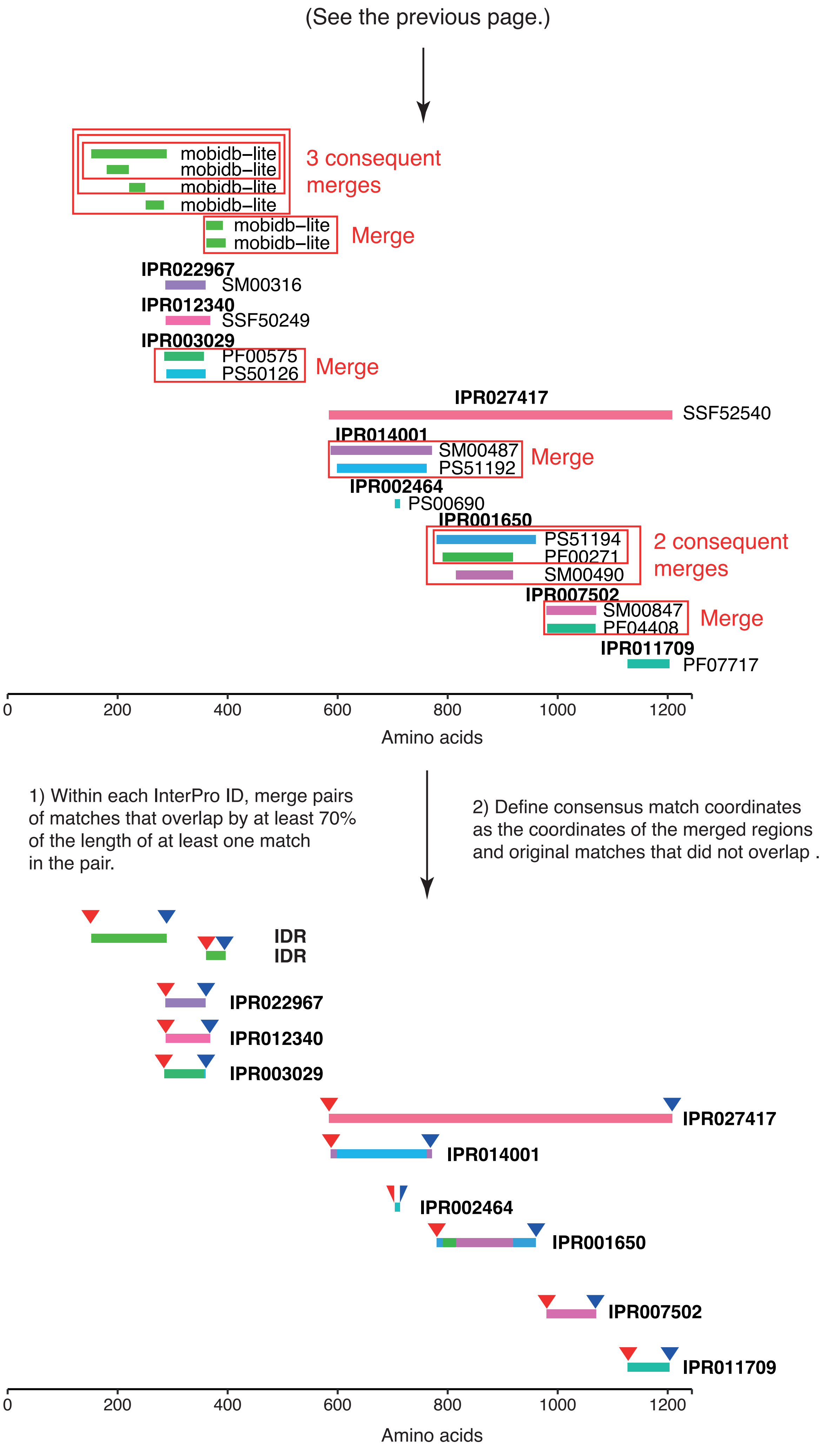
(See the previous page.)
mobidb–lite
mobidb–lite
mobidb–lite
mobidb–lite
3 consequent merges
mobidb–lite
mobidb–lite
Merge
IPR022967
SM00316
IPR012340
SSF50249
IPR003029
PF00575
PS50126
Merge
IPR027417
SSF52540
IPR014001
SM00487
PS51192
Merge
IPR002464
PS00690
IPR001650
PS51194
PF00271
SM00490
2 consequent merges
IPR007502
SM00847
PF04408
Merge
IPR011709
PF07717
0 200 400 600 800 1000 1200
Amino acids
1) Within each InterPro ID, merge pairs of matches that overlap by at least 70% of the length of at least one match in the pair.
2) Define consensus match coordinates as the coordinates of the merged regions and original matches that did not overlap .
IDR
IDR
IPR022967
IPR012340
IPR003029
IPR027417
IPR014001
IPR002464
IPR001650
IPR007502
IPR011709
0 200 400 600 800 1000 1200
Amino acids

G

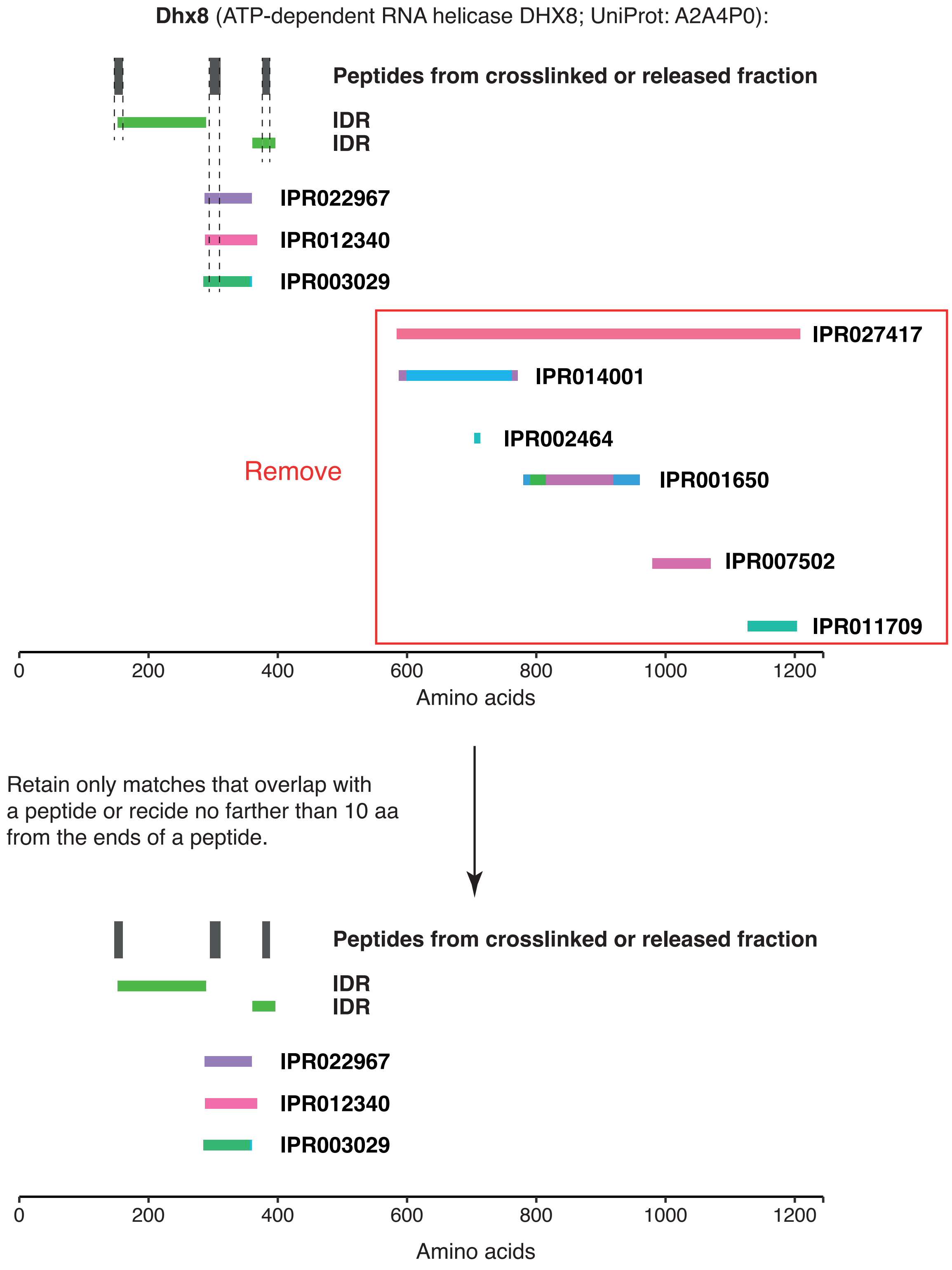

**Dhx8** (ATP-dependent RNA helicase DHX8; UniProt: A2A4P0):
Peptides from crosslinked or released fraction
IDR
IDR
IPR022967
IPR012340
IPR003029
IPR027417
IPR014001
IPR002464
IPR001650
IPR007502
IPR011709
Remove
0
200
400
600
800
1000
1200
Amino acids
Retain only matches that overlap with
a peptide or recide no farther than 10 aa
from the ends of a peptide.
Peptides from crosslinked or released fraction
IDR
IDR
IPR022967
IPR012340
IPR003029
0
200
400
600
800
1000
1200
Amino acids

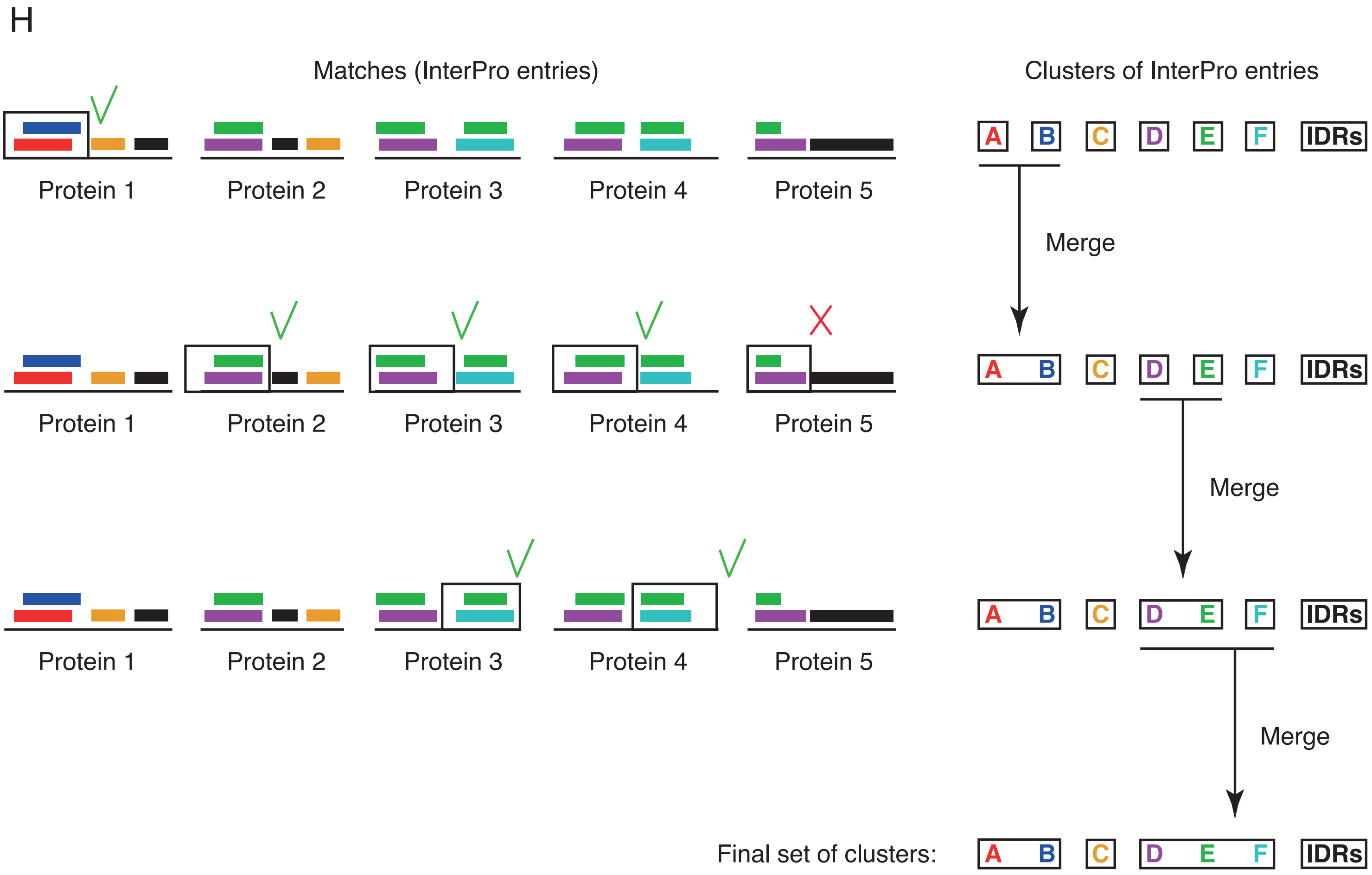
H
Matches (InterPro entries)
Clusters of InterPro entries
Protein 1
Protein 2
Protein 3
Protein 4
Protein 5
A
B
C
D
E
F
IDRs
Merge
Merge
Merge
Final set of clusters:

### Suppl. Figure 5

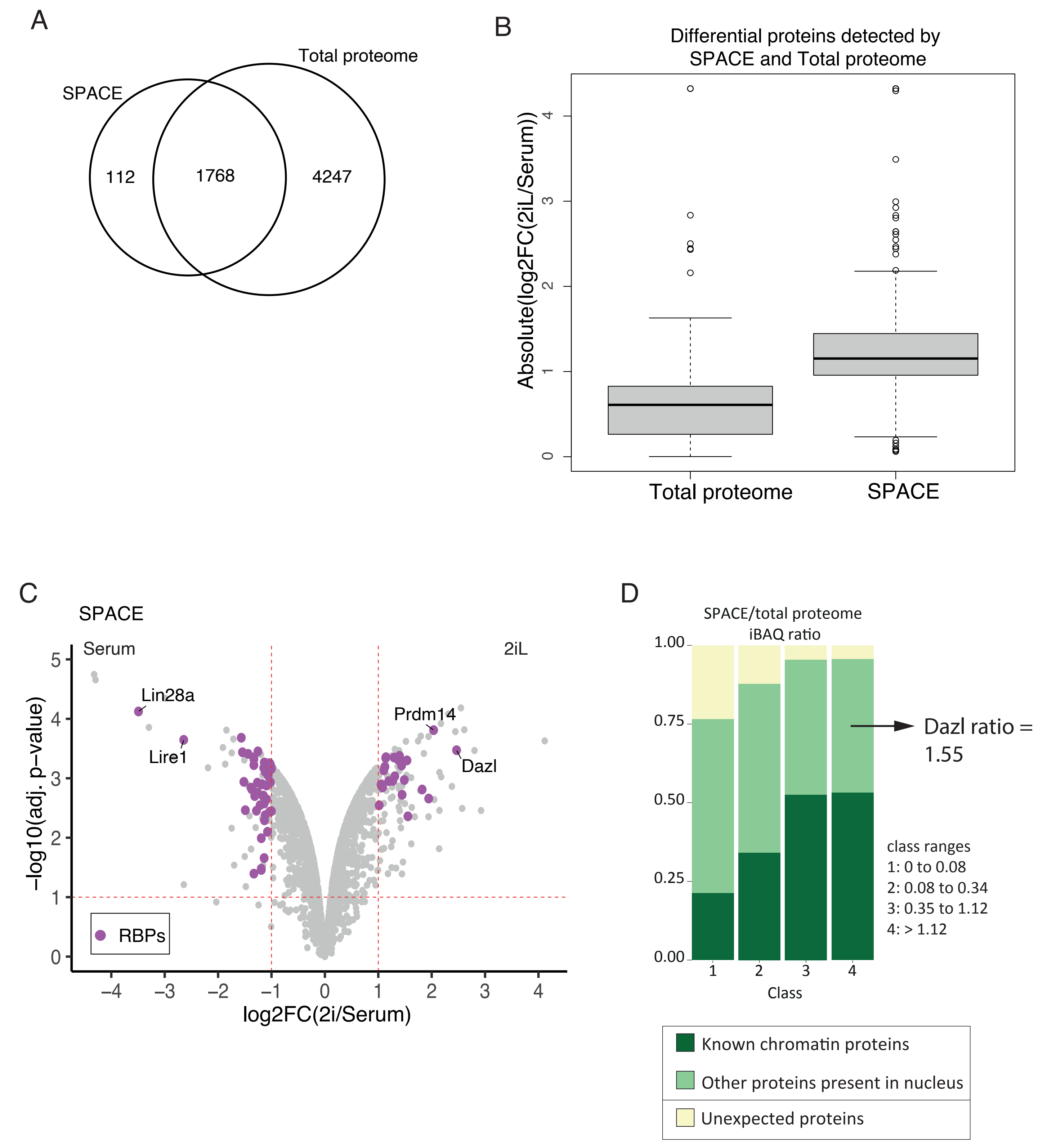

### Suppl. Figure 6

A

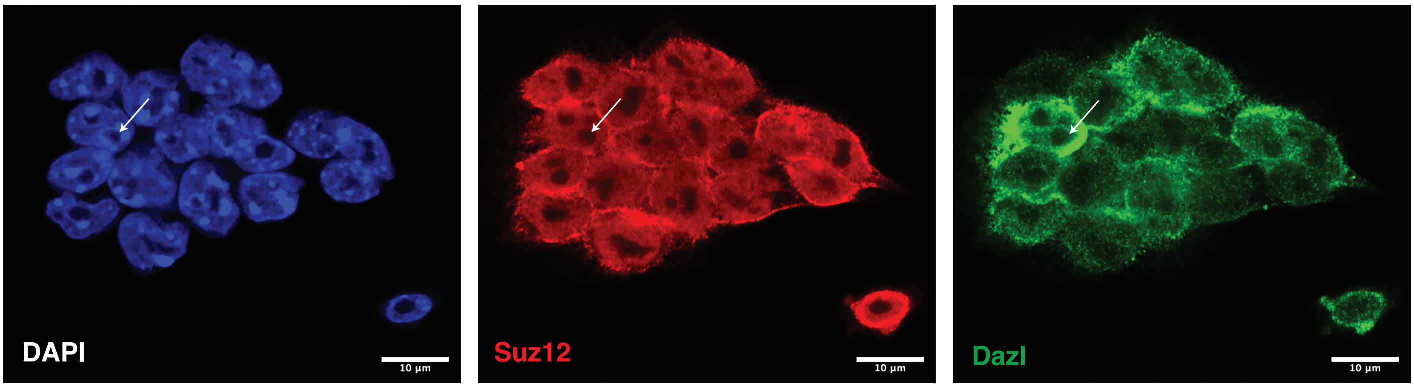


B

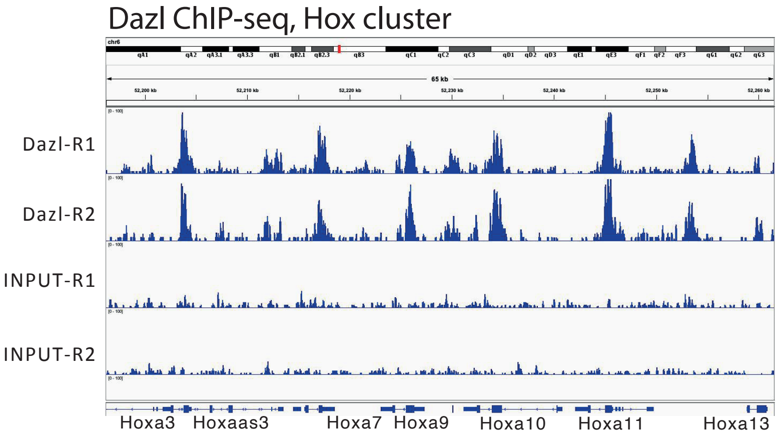


C

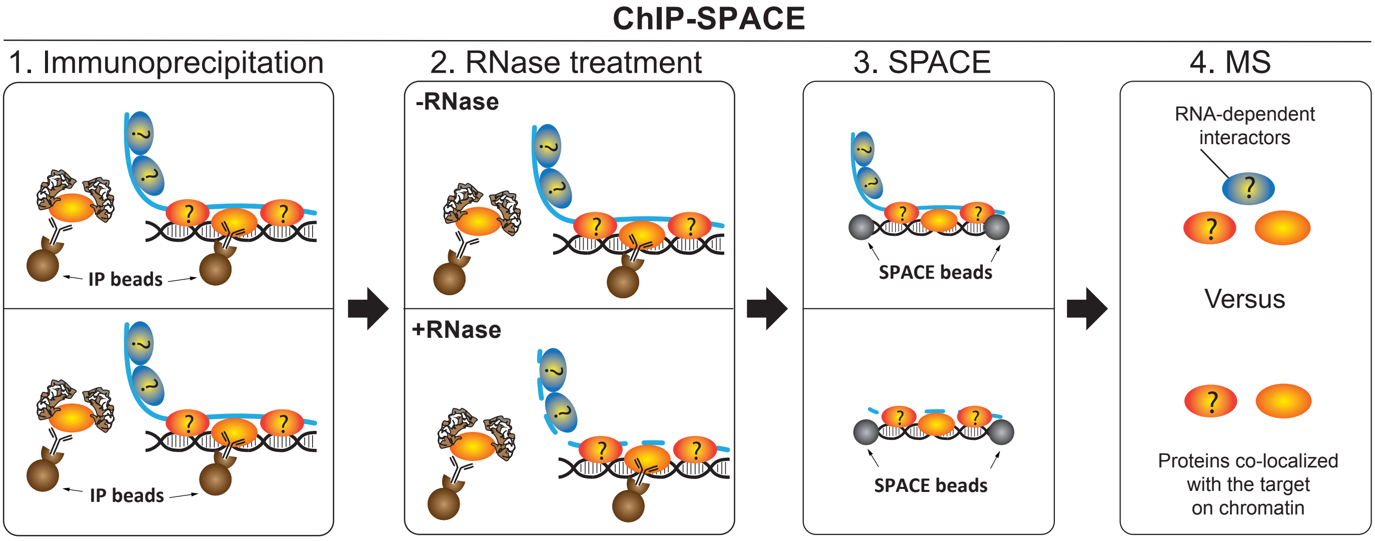
